# Transcriptional responses of *Escherichia coli* during recovery from inorganic or organic mercury exposure

**DOI:** 10.1101/161646

**Authors:** Stephen LaVoie, Anne O. Summers

## Abstract

**Background:** The protean chemical properties of mercury have long made it attractive for diverse applications, but its toxicity requires great care in its use, disposal, and recycling. Mercury occurs in multiple chemical forms, and the molecular basis for the distinct toxicity of its various forms is only partly understood. Global transcriptomics applied over time can reveal how a cell recognizes a toxicant and what cellular subsystems it marshals to repair and recover from the damage. The longitudinal effects on the transcriptome of exponential phase *E. coli* were compared during sub-acute exposure to mercuric chloride (HgCl_2_) or to phenylmercuric acetate (PMA) using RNA-Seq.

**Results:** Differential gene expression revealed common and distinct responses to the mercurials throughout recovery. Cultures exhibited growth stasis immediately after each mercurial exposure but returned to normal growth more quickly after PMA exposure than after HgCl_2_ exposure. Correspondingly, PMA rapidly elicited up-regulation of a large number of genes which continued for 30 min, whereas fewer genes were up-regulated early after HgCl_2_ exposure only some of which overlapped with PMA up-regulated genes. By 60 min gene expression in PMA-exposed cells was almost indistinguishable from unexposed cells, but HgCl_2_ exposed cells still had many differentially expressed genes. Relative expression of energy production and most metabolite uptake pathways declined with both compounds, but nearly all stress response systems were up-regulated by one or the other mercurial during recovery.

**Conclusions:** Sub-acute exposure influenced expression of ~45% of all genes with many distinct responses for each compound, reflecting differential biochemical damage by each mercurial and the corresponding resources available for repair. This study is the first global, high-resolution view of the transcriptional responses to any common toxicant in a prokaryotic model system from exposure to recovery of active growth. The responses provoked by these two mercurials in this model bacterium also provide insights about how higher organisms may respond to these ubiquitous metal toxicants.

## BACKGROUND

The common metallic element mercury (Hg) has no beneficial biological function and its chemical similarities to essential transition metals such as zinc, copper, and iron make it highly toxic to all living systems. Global mercury emissions range from 6500 to 8500 Mg annually with estimates of half [1, 2] and even two-thirds [3] being anthropogenic and the rest from volcanism. Mercury exists in multiple chemical forms that are readily susceptible to abiotic and biotic inter-conversions [4]. Mercury occurs naturally as the insoluble HgS ore (cinnabar), as inorganic complexes of Hg^+2^, Hg^+1^, or 2+ (Hg_2_)^2+^ of varying solubility depending on available ligands, and as organomercurials generated by microbial and anthropogenic processes.

Major sources of chronic mercury exposure in humans include dental amalgam fillings [5, 6], consumption of fish containing methylmercury [7], and artisanal gold mining operations [8]. Organomercurials, like phenylmercury, methylmercury, and merthiolate (ethylmercury) have historically been used in medical, industrial and agricultural applications as antimicrobial or fungicidal agents [9–11]. The toxic effects of mercury exposure in humans are associated with neurological, kidney, liver, gastrointestinal, and developmental disorders [9, 12–15].

Like other common electrophilic toxic metals such as arsenic, cadmium, and lead, there is no single biochemical target for mercury damage. Mercury has a strong affinity for sulfur and selenium [16, 17] and therefore targets the cellular thiol pool, composed of glutathione and cysteine thiol groups of proteins [9] and selenocysteine, a rare but critical amino acid in proteins involved in redox defense and thyroid function [18]. Depletion of the cellular thiol pool and disruption of the cellular membrane potential by mercury can induce oxidative stress and apoptosis pathways in mitochondria [19, 20]. However, there is no evidence that mercury itself undergoes Fenton-type chemistry to generate reactive oxygen species like iron and copper [21].

In earlier work we used global proteomics to identify stable mercury-protein binding sites in growing *E. coli* cells exposed to acute levels of organic or inorganic Hg [22]. We found cysteine sites in several hundred proteins, many highly conserved evolutionarily, that formed stable adducts with one or more of these mercurials, consequently disrupting many cellular processes such as iron homeostasis and the electrolyte balance [23]. Importantly, we found that organic and inorganic mercurials had distinct effects on these cellular processes and distinct protein structural preferences. Although the pathobiology of organic and inorganic mercurials has been known for decades to differ, with methyl- and ethyl-mercury recognized as neurotoxic and inorganic mercury as neurotoxic, nephrotoxic, hepatotoxic, and immunotoxic, no previous studies at that time had assessed the biochemical underpinnings of these distinctions on a global scale in any model system.

Motivated by our proteomics observations and by microarray data from *C. elegans* showing distinct transcriptional single end point response and toxicity for inorganic and organic mercurials [24], we applied RNA-Seq to examine the transcriptional effects of HgCl_2_ and phenylmercuric acetate (PMA) exposure on *E. coli* K-12 MG1655 over time. This is the first study to examine the global transcriptional response to mercury exposure in a microorganism and the only study to compare directly the effects of different compounds over time through recovery. The changes in gene expression were idiosyncratic for each compound, confirming and extending the idea that the cell suffers overlapping but distinct biochemical damage and marshals both distinct and overlapping recovery processes in response to these chemically distinct mercurials. Although our work was in a bacterium, the high evolutionary conservation of many of the genes whose expression we identified as mercury-vulnerable offers insights for the toxicology of mercury compounds in higher organisms.

## METHODS

### Cell cultures

For each biological RNA-Seq replicate *E coli* K-12 MG1655 was subcultured from cryostorage on Luria-Bertani (LB) agar overnight at 37°C. A half-dozen well-isolated colonies were used to inoculate a 20 ml starter culture in Neidhardt MOPS Minimal Medium (NM3) [25] (0.2% final glucose concentration) supplemented with 20 mg/L uracil and 500 μg/L thiamine, which was incubated at 37°C with shaking at 250 rpm overnight (~18 hr). The overnight starter culture was diluted 1:30 to initiate the experimental culture and divided into three 500 ml flasks with 100 ml NM3 in each, which were incubated at 37°C with shaking at 250 rpm. When cultures reached OD_595_ ≈ 0.470 (~ 200 min), two cultures were made 3 μM mercuric chloride (HgCl_2_) or 3 μM phenylmercuric acetate (PMA) and the third was left as an unexposed control. Mercury stocks were prepared fresh for each growth experiment: 10 mM HgCl_2_ (Fisher) in water and 5 mM PMA (Sigma) in 25% dimethyl sulfoxide DMSO (Fisher), which is 2.1 mM or 0.015% v/v final concentration DMSO in culture. These mercurial exposures were chosen from prior pilot experiments to find exposure conditions (OD595, mercurial concentration and sampling times) that displayed a marked decrease in growth rate relative to the unexposed control but allowed subsequent restoration of rapid growth rate (i.e. recovery) within 1 hr (approximately one generation in NM3) after mercurial exposure. Duplicate 1-ml aliquots of each culture were collected at 0 (unexposed control only), 10, 30, 60 min after mercurial exposure and immediately centrifuged at 20,800 x *g* for 3 min at 4°C. Spent medium was aspirated and cell pellets were frozen at −70°C within 5 min after collection. Seven biological replicates were prepared following this protocol and the average variance for all replicates in culture optical density over each 90-min experiment ranged from 0.0019 – 0.0073. The three biological replicates with the lowest variance between growth curves (range from 0.0007 – 0.0017 for all time points) were prepared for RNA-Seq.

### Purification of mRNA

One cell pellet from each condition and sampling time was thawed on ice for all three biological replicates; total RNA was isolated by RNA*snap*™ [26] and stored at - 70°C. DNA contamination was removed by two treatments with Turbo-DNase (Ambion; Life Technologies). RNA concentrations and A_260_/A_280_ ratios were determined using a Nanodrop™ 1000 spectrophotometer (Thermo Scientific). Ribosomal RNA depletion was performed with the Ribo-Zero™ rRNA removal kit for Gram-negative bacteria (Epicentre) and concentrated using RNA Clean and Concentrator– −5 columns (Zymo Research) following the manufacturer’s instructions. Purified mRNA was quantified using the Nanodrop™ and stored at −70°C.

### Library Preparation and Next-generation Sequencing

The quality and quantity of rRNA-depleted RNA was assessed on a 2100 Bioanalyzer RNA pico chip (Agilent Technologies) using the manufacturer’s recommendations. Next-generation sequencing (NGS) libraries were prepared using the Kapa biosystems NGS stranded library prep kit for RNA-Seq with dual indexed Illumina adapters. Library insert size was ~150 bp, as determined by high-sensitivity NGS fragment analysis kit for Fragment Analyzer™ (Advanced Analytical Technologies) using the manufacturer’s instructions. Quantification of each library was done by qPCR and all 30 libraries were pooled in equal concentrations. The library preparation, quality analysis, and pooling were performed by the Georgia Genomics Facility (http://dna.uga.edu). Paired-end (2 x 50 bp) sequencing of the pooled libraries using the Illumina HiSeq 2000 platform was performed by the HudsonAlpha Institute for Biotechnology Genomic Services Laboratory (http://gsl.hudsonalpha.org). See Table S1 for index and filename information for data uploaded to NCBI Gene Expression Omnibus database (http://www.ncbi.nlm.nih.gov/geo/) with accession ID: GSE95575.

### Data Processing and Differential Expression Analysis

Quality control processing of sequence data was performed using Galaxy (https://galaxyproject.org) on the Georgia Advanced Computing Resource Center at the University of Georgia. The FASTX tools in Galaxy (http://hannonlab.cshl.edu/fastx_toolkit) were used for filtering by quality (80% of sequence ≥ quality score of 20), then reads were trimmed at both 5’ and 3’ ends using a window and step size of 1 with quality score ≥ 20. Forward- and reverse-read mate-pairs were assembled and aligned to the *Escherichia coli* MG1655 K-12 genome using Bowtie2 [27]. SAMtools [28] was used to convert Bowtie2 output (.bam file) to SAM format. The number of sequence reads that aligned to features in the annotation file (Escherichia_coli_str_k_12_substr_mg1655.GCA_000005845.2.24.gtf from http://bacteria.ensembl.org) were tabulated from the resulting SAM alignment files using the HTSeq-count program [29] with intersection non-empty mode. Mapped read counts were analyzed for significant differential expression (false discovery rate of ≤ 0.01, fold-change ≥ 2) using the baySeq package in R [30]. All genes that did not meet both the ≤ 1% FDR and ≥ 2 fold-change criteria were indicated as no-change in figures, tables, and text. Within baySeq, two-way comparisons using quantile normalization were made for all three biological replicate transcriptomes over time for HgCl_2_ exposure or PMA exposure versus the unexposed control. We also examined changes over time in the unexposed control culture itself.

## RESULTS

### Effects of sub-acute mercury exposure on growth of MG1655

We defined sub-acute exposure as the concentration of mercury that clearly inhibited growth relative to the unexposed control but allowed cells to resume growth within 1 hour or approximately one generation in this medium (Figure 1a, Figure S1).

**Figure 1:**
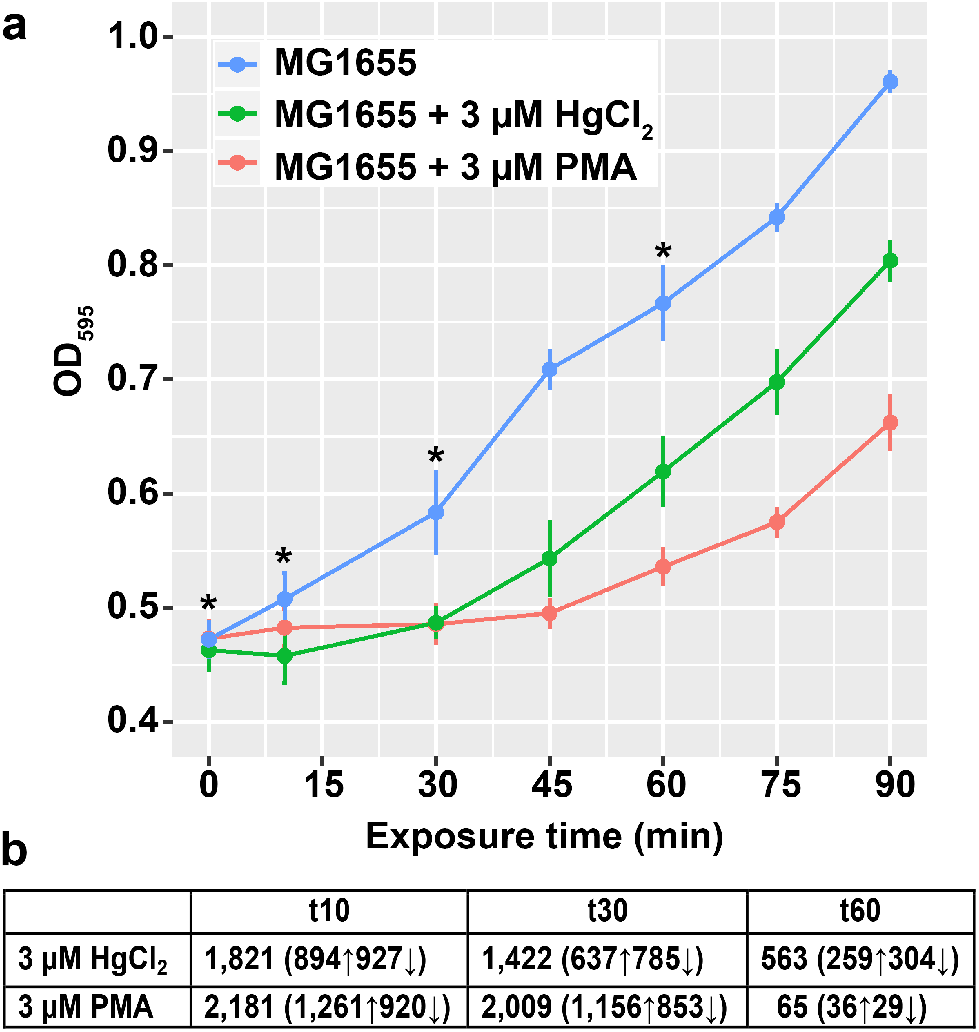
Effects of sub-acute mercury exposure on growth of MG1655. **(a)** *E. coli* K12 MG1655 grown in MOPS minimal medium, unexposed (blue) or exposed to 3 μM HgCl_2_ (red) or 3 μM PMA (green) during mid-log phase. Asterisks indicate sampling times for RNA-seq. Error bars are standard error (SEM) of 3 biological replicates for each culture condition. See Figure S1 for full growth curve. **(b)** Significantly differentially expressed genes (DEG) counts (up-regulated or down-regulated) for HgCl2 and PMA relative to unexposed control culture at each time point.

We chose restoration of growth rate, not stationary phase cell density, as the benchmark for recovery so as not to conflate normal cellular stationary phase “stop growing” signals with mercury-induced “stop growing” signals. Based on pilot experiments the appropriate dose proved to be 3 μM for both mercurials. Exposure to 4 - 5 μM HgCl_2_ prevented growth resumption during 1 hr and the effects of PMA exposure were similar at 3 and 5 μM; exposure to 2.5 μM of either mercurial did not consistently retard growth (data not shown).

Cell-associated Hg (Table S2 and Supporting Information Methods) declined slowly as has been reported previously for low level HgCl_2_ exposure of Hg sensitive cells and was attributed to non-specific endogenous reductants [31–33]. Bound Hg in cells exposed either to HgCl_2_ or to PMA declined similarly from ~50% of total Hg added to culture at 10 min to ~20% at 30 min, after which Hg loss from PMA-exposed cells continued to decline to 11% of input at 60 min. In contrast, cell-associated Hg in cells exposed to HgCl_2_ increased from 24% at 30 to 47% at 60 min. Presently, we have no simple explanation for this unexpected difference in cell-bound Hg in late exponential phase cultures, however it does echo our finding that cultures acutely exposed to 40 μM or 80 μM PMA or HgCl_2_ bound 24% or 208% more Hg(II) than PhHg, respectively [23]. Also notable was a brief drop in the culture optical density immediately after PMA exposure consistent with some cell lysis as has been reported [34]. The lack of apparent lysis after divalent HgCl_2_ exposure may be due to its ability to cross-link cell envelope proteins via their cysteines, which is not possible for monovalent PMA.

### Transcriptome benchmarks

Paired-end libraries averaged over 9.5 million reads and mapped reads provided an average of 143X coverage (Figure S2). The sequencing data were of high quality, requiring removal of only 11% as low-quality reads. Of the high-quality reads, 97% of reads mate-paired, 99.4% of paired reads mapped to the genome and 82% of reads mapped to an annotated genome feature on average from all libraries.

Overall 89% of annotated mapped reads were to coding regions (CDS) based on raw un-normalized read counts per gene output from HTseq-count program [29](Figure S3, Table S3, Table S4). Pearson correlations of raw read counts confirmed that no strong biases were introduced in biological replicates for each condition (Figures S4-S6). That dispersion is slightly greater in both mercury exposure conditions than in unexposed cultures, especially at later time points, is consistent with perturbations of multiple cellular processes.

Mapped reads to rRNA constituted only 0.3% of total reads (std. dev. = 0.425) on average for all libraries (Table S3) consistent with effective Ribo-Zero™ rRNA removal. In the unexposed culture non-coding RNA’s (ncRNA) were 4% of total reads over all time points, but their percentage increased in mercury exposure conditions indicating greater differential expression of some ncRNA genes (details below). The very abundant tmRNA (*ssrA*) needed for rescuing stalled ribosomes [35] was 6% of total reads in the unexposed condition and although this percentage increased for mercury exposure conditions, the tmRNA gene (*ssrA*) was not differentially expressed under any condition. Pseudogenes accounted for less than 1% of total reads, but up to 35% (HgCl_2_) and 13% (PMA) of them were significantly up-regulated. The tRNA’s were less than 1% of total reads because the library preparation method we used was not optimized for such small RNAs. However, approximately 35% of these observed tRNA genes were significantly down-regulated during the first 30 min after exposure to either mercurial.

### HgCl_2_ and PMA transcriptional responses are not the same

We expected that significantly differentially expressed genes (DEGs) in the mercury exposure conditions (compared to the unexposed cells) would change over time as the cells transitioned from initial growth stasis back into a normal growth rate. We also expected that some DEG responses would be similar because both mercurials are thiophilic and will bind to the cellular thiol redox buffer, glutathione (GSH), and to protein cysteine residues. However, since there are physiological differences and protein site preferences for each compound in acute Hg(II)- or PMA-exposure [23 and Zink et. al. in preparation] we aimed here at a low exposure using a longitudinal protocol to discern more subtle distinctions between these mercurials as the cells experienced stasis and then recovered their growth rate. In the following sections, we first describe the bulk measures of gene expression over time and then describe differences in specific functional pathways.

### Differentially expressed genes: the view from 30,000 feet

#### (a) Differentially expressed genes (DEG) for each condition and time point

DEGs were determined by pairwise comparisons of mercury exposure conditions relative to the unexposed culture at the corresponding time point (Figure 1b and Table S5). Ten minutes after exposure, expression of 41 % or 49% of the 4,472 non-rRNA genes changed significantly for HgCl_2_ or PMA-exposed cells, respectively (Figure 1b).

At 30 min with growth still arrested, 32% of genes in the HgCl_2_-exposed cells were differentially expressed (Figure 1b), (Figure 1a, red). In contrast, PMA-exposed cells at 30 min began to recover their prior growth rate (Figure 1a, green), but 45% of their genes remained differentially expressed (Figure 1b). By 60 min, the PMA-exposed cells were growing at nearly their pre-exposure rate and only 1.5% of genes were differentially expressed, whereas the HgCl_2_-exposed cells were still growing more slowly than pre-exposure with 13% of their genes still differentially expressed compared to the unexposed cells (Figure 1b).

#### (b) Shared and unique genes at each time point for each exposure

The total distinct DEGs across all time points was slightly lower for HgCl_2_ (2,327) than for PMA (2,541) exposure (Figure 1b and Figure S7). More striking were the differences in DEGs at each time point; PMA-exposed cells modulated 20% more genes at 10 min (2,181 vs 1,821) and 40% more at 30 min (2,007 vs 1,422) than HgCl_2_ exposed cells. This trend completely reversed by 60 min when DEGs declined in both exposure conditions but HgCl_2_ exposed cells were still modulating ~9-fold more genes (563) than PMA-exposed cells (65), consistent with the latter recovering normal growth sooner (Figure 1a). Also notable is the carryover of DEGs from one time point to the next where HgCl_2_ exposed cells have 1,001 DEGs in common at 10 and 30 min after exposure but that number is 65% greater in PMA-exposed cells (1,650) (Figure S7). However, HgCl_2_ exposed cells have 52% more DEGs that are unique at 10 min compared to PMA-exposed cells (804 vs 529), i.e. more HgCl_2_ provoked DEGs occur sooner after exposure than later. HgCl2 exposure also yields more DEGs that occur at all time points compared to PMA (Figure S7) since there are very few DEGs at 60 min for PMA exposure.

#### (c) Up-regulated vs. down-regulated genes for each exposure

Sorting expression simply into genes up-regulated or down-regulated by HgCl_2_ or PMA at each time point (Figure 2) revealed additional quantitative distinctions between them. For up-regulated genes, PMA-provoked more unique DEGs than DEGs in common with HgCl_2_ exposure at all time points, in contrast to HgCl_2_ which had more DEGs in common than unique at all but the last time point. This trend was not continued for down-regulated genes where more genes were in common for both compounds than unique at all time points.

**Figure 2:**
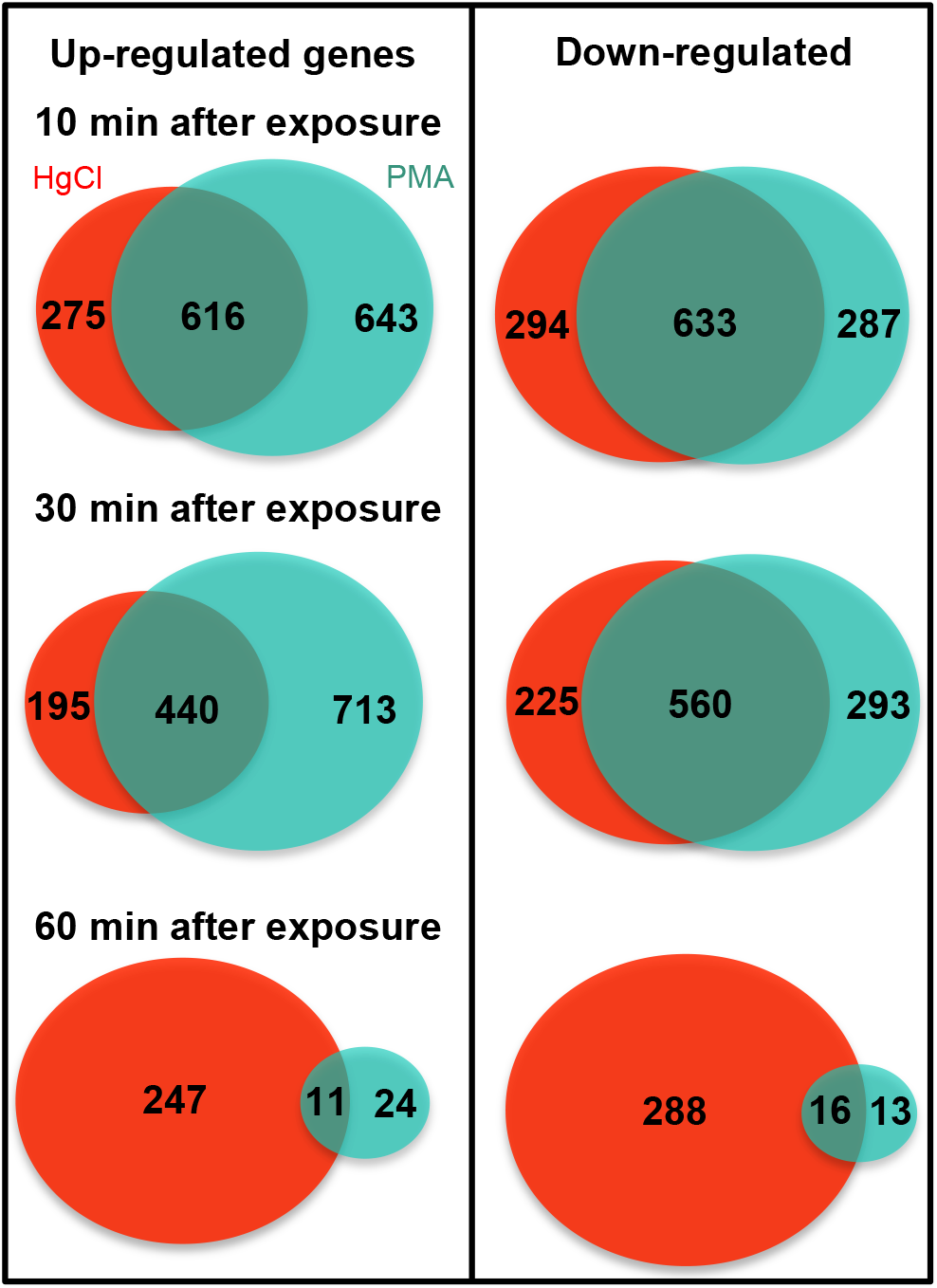
Overlap between differentially expressed genes at each sampling time. The 3 μM HgCl_2_ exposure is in red and the 3 μM PMA exposure in green. Ovals are to scale only at each time point, but not between between time points in a panel nor between left and right panels.

Thus, early in exposure the cell reduced expression of a similar number of genes for both mercurials, but up-regulated expression of many more genes in response to PMA than to HgCl_2_, and these distinct trends persisted to the middle time point. By 60 min, gene expression of PMA-exposed cells closely resembled that of unexposed cells, but HgCl_2_ exposed cells still have many up- and down-DEGs, consistent with slower growth rate recovery by HgCl_2_ exposed compared to PMA-exposed cells (Figure 1a).

#### (d) Differentially expressed genes grouped by functional category

To sort our observations from a different perspective, the DEGs for each condition were grouped by Clusters of Orthologous Groups (COGs) to identify expression differences based on gene functions (Figure 3 and Table S6). For most COGs, both mercurials elicited their strongest responses, up- or down-regulated, within the first 10 min of exposure (Figure 3); in most cases, the responses were of similar magnitude. At 30 min, the PMA (green) responses remained nearly the same, up- or down-regulated, but HgCl_2_ provoked responses (red) generally diminished, often sharply. By 60 min, expression in PMA-exposed cells was barely distinguishable from the unexposed cells in all COG categories, whereas HgCl_2_ exposed cells had notable differential expression in most COG categories.

**Figure 3:**
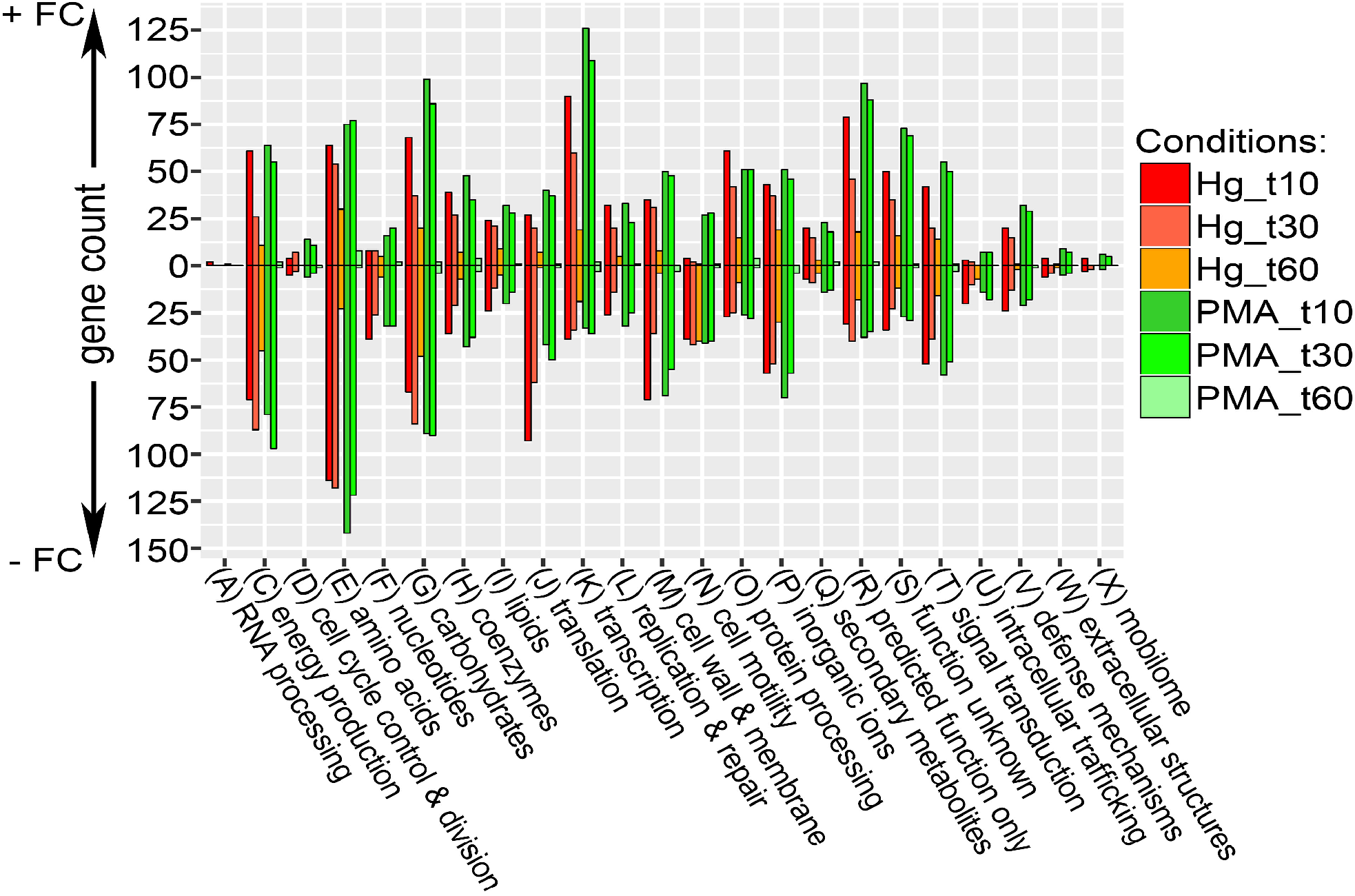
Counts of differentially expressed genes for each condition grouped by COG functional category. Genes with a log2 fold-change ≥ 1 for each condition were grouped by COG group. Positive counts represent observed up-regulated genes and negative counts represent observed down-regulated genes. COG code, number of proteins encoded by genome and category description: (A, 2) RNA processing and modification; (C, 284) Energy production and conversion; (D, 39) Cell cycle control, cell division, chromosome partitioning; (E, 355) Amino acid transport and metabolism; (F, 107) Nucleotide transport and metabolism; (G, 381) Carbohydrate transport and metabolism; (H, 179) Coenzyme transport and metabolism; (I, 121) Lipid transport and metabolism; (J, 236) Translation, ribosomal structure and biogenesis; (K, 294) Transcription; (L, 139) Replication, recombination and repair; (M, 242) Cell wall, membrane and envelope biogenesis; (N, 102) Cell motility; (O, 156) Post-translational modification, protein turnover, chaperones; (P, 223) Inorganic ion transport and metabolism; (Q, 68) Secondary metabolites biosynthesis, transport and catabolism; (R, 261) General function prediction only; (S, 203) Function unknown; (T, 191) Signal transduction mechanisms; (U, 50) Intracellular trafficking, secretion, and vesicular transport; (V, 91) Defense mechanisms; (W, 31) Extracellular structures; (X, 60) Mobilome, prophages, transposons.

The four well-defined COG categories with the most DEGs were energy production (C), amino acid metabolism (E), carbohydrate metabolism (G), and transcription (K), which are also categories with a large number of genes (284, 355, 381, and 294 respectively) in *E. coli* (Figure 3 legend). COG categories R and S encoding poorly defined (261) or non-defined (203) genes were also represented proportionally. These data suggest both mercurials broadly affect most metabolic categories, albeit to different degrees and at different rates. However, four well-defined COG categories have strikingly different responses to HgCl_2_ and PMA. COG categories for nucleotide metabolism (F), translation (J), motility (N), and intracellular trafficking (U) have many fewer up-regulated genes in HgCl_2_ exposed cells than in PMA-exposed cells. There are relatively few DEGs involved in cell division (D), extracellular structures (W), and mobile genetic elements (X) and genes within these categories responded similarly to both mercurials. Thus, grouping DEGs by COGs illuminates the broad functional differences and similarities between HgCl_2_ and PMA exposure. Moreover, this view makes clear that DEGs occur in all functional categories for both compounds, but still display distinct differences in global transcriptional response to each compound. We provide gene-level detail on several of these functional groups below. Note that the COG database only includes functional annotations for 3,398 of *E. coli’s* 4,497 genes, but functional categories discussed below in more detail are not limited to COG-annotated genes.

A heat map of the DEGs log-fold changes (Figure 4) provides a more granular look at all DEGs for both compounds across all time points. The heat map, using Ward’s minimum variance clustering method [36], shows considerable uniformity of up- and down-regulated expression during the 30 min after PMA exposure. In contrast, although HgCl_2_ exposed cells grossly shared many DEGs with PMA-exposed cells (Figure 2), the heat map reveals a more variegated response to Hg(II) in which different genes are up- or down-regulated at all time points, in contrast to the relatively consistent response of PMA during recovery. Overall, 34% of DEGs are unique to only one mercurial and ~3% of all DEGs had an opposite response to each compound (Table S5). The most dramatic differences were at 60 min when HgCl_2_ exposed cells were still modulating many genes but PMA-exposed cells had only minor differences with unexposed cells, consistent with their faster recovery of normal growth (Figure 1a). We dissect some of these differences in function-specific heat maps below.

**Figure 4:**
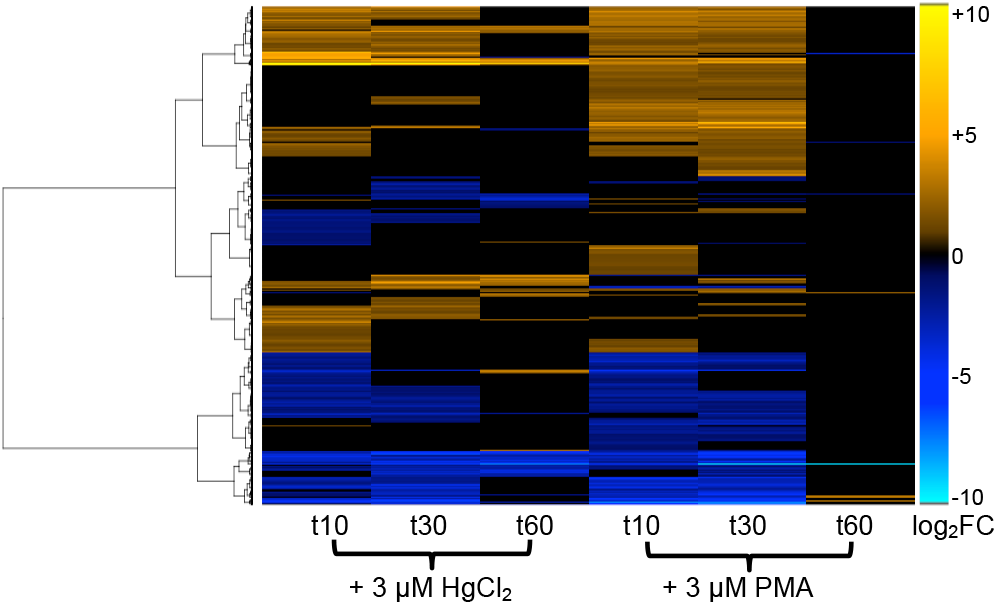
Differentially expressed genes at each RNA sampling time. Heat map of all genes that were differentially expressed in at least one mercury exposure condition (n = 3,149). Genes were clustered by row using Ward’s minimum variance method [36] with non-squared log_2_ fold-change input values.

We also used STRING (version 10.0) [37, 38] for unsupervised network analysis to identify gene clusters that were up-regulated in response to each compound (Figure S8 and Table S7). We focused on up-regulated genes on the working assumption that they are more likely to contribute to recovery than genes whose expression is turned down. Gene clusters were generated by STRING based on organism specific data mining to identify genes with a functional association, such as a common biological purpose, location within the same operon, or shared regulatory mechanism. Note that this network algorithm does not consider fold-change intensity of response; it enumerates only whether an up-regulated gene is present at a given time point. The up-regulated DEGs (nodes) of HgCl_2_ exposed cells formed several tight clusters encompassing 16 gene-ontology functions (GOFs) at 10 min, nine GOFs at 30 min, and seven GOFs at 60 min (Figure S8 and Table S7). In contrast, although there were more nodes for PMA-exposure at 10 and 30 min, there were fewer edges yielding no well-defined clusters at 10 min and only two GOFs at 30 min. This network analysis suggests that, although PMA provokes more DEGs than Hg(II) does, there is less functional congruence between the genes involved in the response to PMA. Specific gene and function changes are discussed further in the next section.

Lastly, as a control for using RNA-Seq in a longitudinal experiment, we observed DEGs at sequential time points in the unexposed control culture (Figure S9 and Table S8). As expected, changes were gradual over time with no more than 5% of the genome being differentially expressed from one time point to the next. At the 60 min time point, as the cells approached stationary phase 815 genes were differentially expressed compared to mid-log (time 0). Sorting these DEGs by COGs (Figure S9) and by STRING network analysis (Figure S10 and Table S7) showed, as expected, many DEGs were consistent with normal transitioning from mid-log to late-log phase [39, 40].

### Higher resolution view of expression differences in specific functional groups during recovery from exposure to HgCl_2_ or PMA

Taking the perspective that a toxicant is a kind of signaling molecule, we considered differences in gene expression for the two Hg compounds to reflect how the cell senses the biochemically distinct damage produced by these two metallo-electrophiles as manifest by what tools the cell calls upon to restore its viability. A quick snapshot of the great extent of these compound-specific differences can be seen in the genes with a >20-fold increase in differential expression after HgCl_2_ exposure (Table 1) or PMA exposure (Table 2). Here we emphasize up-regulated genes on the working assumption they could contribute directly or indirectly to repairing damage caused by mercurial exposure. The 25 genes highly up-regulated by HgCl_2_ (Table 1) are involved in altering the cell surface, oxidative stress response and repair, protein chaperones, metals homeostasis, and ribonucleotide reductase. The vestigial prophage genes likely play no rescue role for the cell and were simply activated by generalized stress responses and have large fold-change values due to repressed expression under normal growth conditions. The corresponding PMA response echoed only 8 of these 25 HgCl_2_ high-responders, and notably did not include the vestigial phage genes.

**Table 1:** Genes with ≥ 20 fold-change for at least two time points after HgCl_2_ exposure (n = 25).

| Gene ID | Gene Name | Product Description | Hg t10 | Hg t30 | Hg t60 | PMA t10 | PMA t30 | PMA t60 |
| --- | --- | --- | --- | --- | --- | --- | --- | --- |
| b1112 | **bhsA** | Biofilm, cell surface and signaling defects, YhcN family | **1949** | **563** | **94** | **46** | **21** | n.s. |
| b4458 | oxyS | OxyS sRNA activates genes that detoxify oxidative damage | **1524** | **1292** | **150** | 10 | 6 | n.s. |
| b4249 | bdcA | c-di-GMP-binding biofilm dispersal mediator protein | **596** | **79** | 7 | 16 | 2 | n.s. |
| b3686 | **ibpB** | Chaperone, heat-inducible protein of HSP20 family | **402** | **124** | **281** | **37** | **211** | 9 |
| b0812 | dps | Stress-induced Fe-binding and storage protein | **339** | **52** | n.s. | 4 | 2 | n.s. |
| b4248 | yjgH | Putative reactive intermediate deaminase, UPF0076 family | **296** | **24** | 3 | 6 | 2 | n.s. |
| b0849 | grxA | Glutaredoxin 1 | **246** | **33** | n.s. | 14 | 4 | n.s. |
| b1144 | ymfJ | Function unknown, e14 prophage | **185** | **39** | n.s. | 3 | 3 | n.s. |
| b2582 | **trxC** | Thioredoxin 2, zinc-binding; Trx2 | **134** | **50** | 7 | **24** | **24** | n.s. |
| b1147 | ymfL* | Function unknown, e14 prophage | **95** | **29** | n.s. | n.s. | 3 | n.s. |
| b1146 | croE* | Cro-like repressor, e14 prophage | **90** | **40** | n.s. | n.s. | n.s. | n.s. |
| b3687 | **ibpA** | Chaperone, heat-inducible protein of HSP20 family | **86** | 19 | **41** | 10 | **28** | 4 |
| b2531 | iscR | Transcriptional repressor for isc operon; contains Fe-S cluster | **62** | **48** | 9 | 12 | 18 | n.s. |
| b1148 | ymfM* | Function unknown, e14 prophage | **49** | **21** | n.s. | n.s. | n.s. | n.s. |
| b4599 | yneM | function unknown, membrane-associated; regulated by PhoPQ | **38** | **36** | n.s. | 3 | 3 | n.s. |
| b2673 | nrdH | NrdH-redoxin reducing oxidized NrdEF | **35** | **127** | **21** | 7 | 14 | n.s. |
| b4030 | **psiE** | Pho regulon, regulated by phoB and cAMP | **35** | **64** | 17 | **38** | **59** | n.s. |
| b1684 | sufA | Scaffold protein for assembly of iron-sulfur clusters | **32** | **39** | 5 | n.s. | 3 | n.s. |
| b1748 | **astC** | Succinylornithine transaminase; carbon starvation protein | **30** | **31** | **25** | **34** | **36** | n.s. |
| b4663 | **azuC** | Function unknown; membrane-associated | **27** | **25** | **36** | **41** | **49** | n.s. |
| b0484 | copA | Copper-, silver-translocating P-type ATPase efflux pump | **25** | **20** | n.s. | 5 | n.s. | -10 |
| b2674 | nrdI | Flavodoxin required for NrdEF cluster assembly | **21** | **73** | **22** | 4 | 7 | n.s. |
| b1747 | astA* | Arginine succinyltransferase, arginine catabolism | 10 | **20** | **29** | 9 | 14 | n.s. |
| b1020 | **phoH** | ATP-binding protein, function unknown | 8 | **71** | **221** | **61** | **365** | n.s. |
| b4002 | zraP | Zn-dependent periplasmic chaperone | 3 | **27** | **43** | n.s. | n.s. | -3 |

Table is sorted by Hg at 10 min column. n.s. = not significantly different from unexposed culture and boldface highlights actual values ≥ 20-fold. Gene names in boldface have a $\geq 20$ differential expression response to both Hg and PMA in at least one time point (n = 8). Genes marked with an asterisk (\*) had fewer than 11 average read counts for at least 2 time points in the unexposed condition and were operationally defined as being “off” in that condition. Such genes constituted, the lowest 15th percentile of all genes, based on raw read counts normalized across all libraries and across all biological replicates.

**Table 2:**
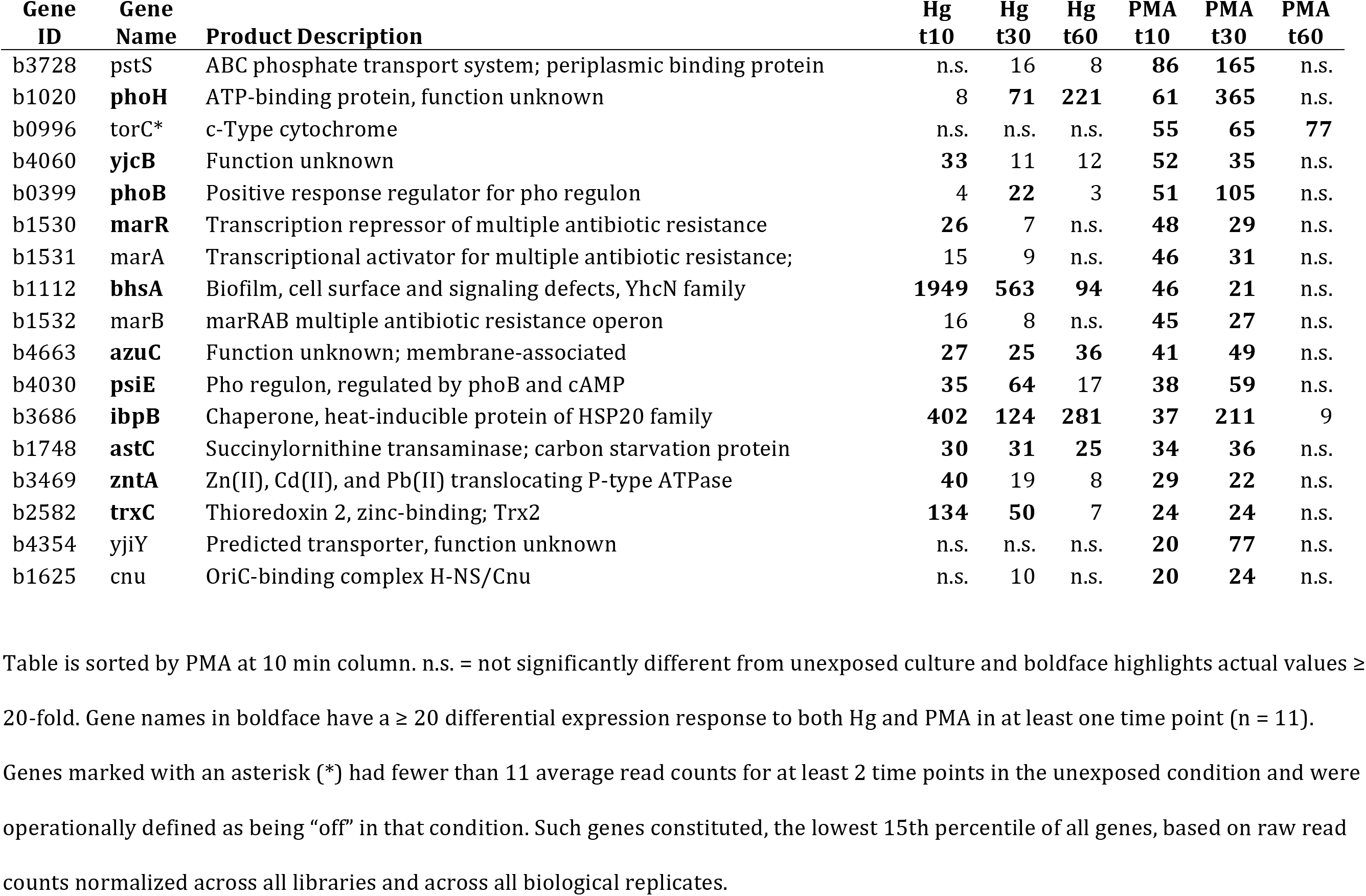
Genes with ≥ 20 fold-change in at least two time points for PMA exposure (n= 17).

| Gene ID | Gene Name | Product Description | Hg t10 | Hg t30 | Hg t60 | PMA t10 | PMA t30 | PMA t60 |
| --- | --- | --- | --- | --- | --- | --- | --- | --- |
| b3728 | pstS | ABC phosphate transport system; periplasmic binding protein | n.s. | 16 | 8 | **86** | **165** | n.s. |
| b1020 | **phoH** | ATP-binding protein, function unknown | 8 | **71** | **221** | **61** | **365** | n.s. |
| b0996 | torC* | c-Type cytochrome | n.s. | n.s. | n.s. | **55** | **65** | **77** |
| b4060 | **yjcB** | Function unknown | **33** | 11 | 12 | **52** | **35** | n.s. |
| b0399 | **phoB** | Positive response regulator for pho regulon | 4 | **22** | 3 | **51** | **105** | n.s. |
| b1530 | **marR** | Transcription repressor of multiple antibiotic resistance | **26** | 7 | n.s. | **48** | **29** | n.s. |
| b1531 | marA | Transcriptional activator for multiple antibiotic resistance; | 15 | 9 | n.s. | **46** | **31** | n.s. |
| b1112 | **bhsA** | Biofilm, cell surface and signaling defects, YhcN family | **1949** | **563** | **94** | **46** | **21** | n.s. |
| b1532 | marB | marRAB multiple antibiotic resistance operon | 16 | 8 | n.s. | **45** | **27** | n.s. |
| b4663 | **azuC** | Function unknown; membrane-associated | **27** | **25** | **36** | **41** | **49** | n.s. |
| b4030 | **psiE** | Pho regulon, regulated by phoB and cAMP | **35** | **64** | 17 | **38** | **59** | n.s. |
| b3686 | **ibpB** | Chaperone, heat-inducible protein of HSP20 family | **402** | **124** | **281** | **37** | **211** | 9 |
| b1748 | **astC** | Succinylornithine transaminase; carbon starvation protein | **30** | **31** | **25** | **34** | **36** | n.s. |
| b3469 | **zntA** | Zn(II), Cd(II), and Pb(II) translocating P-type ATPase | **40** | 19 | 8 | **29** | **22** | n.s. |
| b2582 | **trxC** | Thioredoxin 2, zinc-binding; Trx2 | **134** | **50** | 7 | **24** | **24** | n.s. |
| b4354 | yjiY | Predicted transporter, function unknown | n.s. | n.s. | n.s. | **20** | **77** | n.s. |
| b1625 | cnu | OriC-binding complex H-NS/Cnu | n.s. | 10 | n.s. | **20** | **24** | n.s. |

Table is sorted by PMA at 10 min column. n.s. = not significantly different from unexposed culture and boldface highlights actual values ≥
20-fold. Gene names in boldface have a $\geq 20$ differential expression response to both Hg and PMA in at least one time point (n = 11).
Genes marked with an asterisk (\*) had fewer than 11 average read counts for at least 2 time points in the unexposed condition and were
operationally defined as being “off” in that condition. Such genes constituted, the lowest 15th percentile of all genes, based on raw read
counts normalized across all libraries and across all biological replicates.

For PMA, the highly up-regulated genes are a distinct contrast to those for HgCl_2_. First, the maximum amplitude of the PMA-provoked differential expression is generally much less than for Hg(II)-provoked high differential expression (Table 1), which could reflect the lower uptake of PMA. Secondly, while 11 of the 17 PMA-provoked genes were also on the HgCl_2_ highly differential expression list, ion transport and antibiotic resistance loci were more prominent with PMA and prophage genes were absent.

These two snapshot tables make the points that both mercurials generate broad, but idiosyncratic, cellular responses. To place these “tips of many icebergs” in their larger cellular context, we used heat maps and tables of subsets of functionally related genes to discuss the differential effects of HgCl_2_ and PMA on twelve canonical cellular systems in the following sections.

## i. INFORMATIONAL MACROMOLECULES

### (a) DNA replication, recombination and repair

Of the 24 genes for initiation and maintenance, and termination of chromosome replication, there were more genes down-regulated (8) than up-regulated (3) in response to HgCl_2_ and an equal number up- or down-regulated genes (7) in response to PMA (Table S9). Of the 14 genes encoding the replicative polymerase holoenzyme, four genes capable of translesion synthesis (*polB, dinB, umuCD*) were up-regulated more by HgCl_2_ exposure (Table S9), suggesting a greater degree of direct or indirect DNA damage by HgCl_2_ exposure. Of the 45 genes for repair and recombination proteins the transcriptional response to each mercurial was very similar (11 up-regulated and 16 down-regulated for HgCl_2_; 9 up-regulated and 18 down-regulated for PMA). But there were repair genes unique to each compound: *xthA*, *uvrAB*, *mutM*, and *recN* were only up-regulated by HgCl_2_; and *mutH* and *mutY* were only up-regulated by PMA.

The *recA*, *recN*, and *xthA* DNA repair genes were the most highly up-regulated (≥10 fold) in response only to HgCl_2_. The *recA* gene, induced by double-strand DNA breaks, serves multiple roles in DNA repair [41, 42]. Curiously, expression of *recBCD*, which is needed for break repair, either did not change or declined compared to unexposed cells for both mercurials. Expression of several genes involved in repair (*recG, nth, hsdS*, and *mcrC*) were down-regulated by both compounds, but with larger negative fold-changes for PMA than HgCl_2_. Thus, the cells responded quickly to both mercurials, but some distinct responses suggest these two compounds directly or indirectly yield different kinds of DNA damage.

### (b) Transcription

Of the core RNA polymerase (RNAP) genes only PMA-exposure increased expression of a single gene *rpoZ* (ω subunit), but expression decreased in the remaining *rpoABC* core genes (Table S10). HgCl_2_ exposure did not change expression of any RNAP core genes except for a transient 3-fold drop in *rpoA* at 30 min. Only one of the five termination factors, the Rho-directed anti-terminator, *rof*, increased and did so for both mercurials with PMA again provoking a greater response.

Genes for three sigma factors displayed increased expression upon exposure to either mercurial, with *rpoH* (heat shock sigma factor) and *rpoS* (stationary phase and stress response sigma factor) increasing more following PMA-exposure and *rpoD* (housekeeping sigma factor) only increasing after HgCl_2_ exposure. The effects of HgCl_2_ or PMA exposure on the regulation of genes within each regulon controlled by *E. coli’s* seven sigma factors are tabulated in Table S11. Many genes are modulated differentially by HgCl_2_ or PMA-exposure, but no single sigma factor is uniquely responsible for increases or decreases in responses to these two compounds.

Many of the 203 transcriptional regulators annotated in the RegulonDB (Table 3) [43, 44] and the 1,723 genes they control were expressed differently with the two mercurials (Table S12). PMA provoked up-regulation of more transcription factor genes at 10 and 30 min than HgCl_2_ exposure, but slightly fewer down-regulated regulators (Table 3). Of all COG categories, transcription had the most up-regulated genes for both mercurials (Figure 3 and Table S6). PMA up-regulated ~40% more transcription related genes at 10 min and ~80% more genes at 30 min than HgCl_2_. Six activators (*mhpR*, *glcC*, *gadX*, *soxS*, *mlrA*, *phoB*) and three repressors (*mcbR*, *iscR*, *betI*) were up-regulated at all times for HgCl_2_, but *gadX* was the only activator gene up-regulated at all times for both mercurials (Table S12). GadX is part of the RpoS regulon [45] and activates the acid resistance system and multidrug efflux [46, 47]. Details of transcription factors and their regulons are provided in Table S12.

**Table 3:** Changes in transcription factor gene expression. The sum of transcription factor genes either up-regulated or down-regulated is shown with the percentage of the total transcription factor genes in parenthesis; percents do not total 100 because genes with no change compared to unexposed cells are not tabulated here. See details in Table S12.

| Transcription Factors (n = 203) |  |  |  |  |  |  |
| --- | --- | --- | --- | --- | --- | --- |
|  | Hg_t10 | Hg_t30 | Hg_t60 | PMA_t10 | PMA_t30 | PMA_t60 |
| up | 63 (31) | 46 (23) | 14 (7) | 86 (42) | 78 (38) | 1 (0.5) |
| down | 28 (14) | 28 (14) | 13 (6) | 22 (11) | 23 (11) | 2 (1) |

Lastly, *E. coli* has 65 currently annotated (ASM584v2), small non-coding RNAs. Although our RNA purification and library preparation methods were not optimized for their enrichment, we observed differential expression for a number of them (Table S5, feature type “ncRNA”). ncRNAs up-regulated for both mercurials are involved in regulation of acid resistance (*gadY*), oxidative stress (*oxyS*), and multiple transporters (*gcvB* and *sgrS*). In contrast, adhesion and motility (*cyaR*), and anaerobic metabolism shift (*fnrS*) were down-regulated by both compounds.

### (c) Translation

Upon HgCl_2_ exposure 83% and 74% of ribosomal proteins (r-proteins) were down-regulated at 10 and 30 min, respectively, versus only 4% and 41% for PMA at the corresponding times (Figures 5 and S9; data for all functional group heat maps are shown in Table S13 and Table S14). Transcription of r-proteins is repressed directly by binding of the nutritional stress-induced nucleotide ppGpp and DksA protein to RNAP [48]. The ppGpp synthase genes, *spoT* and *relA*, were down-regulated or unchanged, but expression of *dksA* was up-regulated for both HgCl_2_ and PMA exposure. R-protein expression can also be inhibited by excess r-proteins binding to and inhibiting translation of their own mRNAs [49, 50].

**Figure 5:**
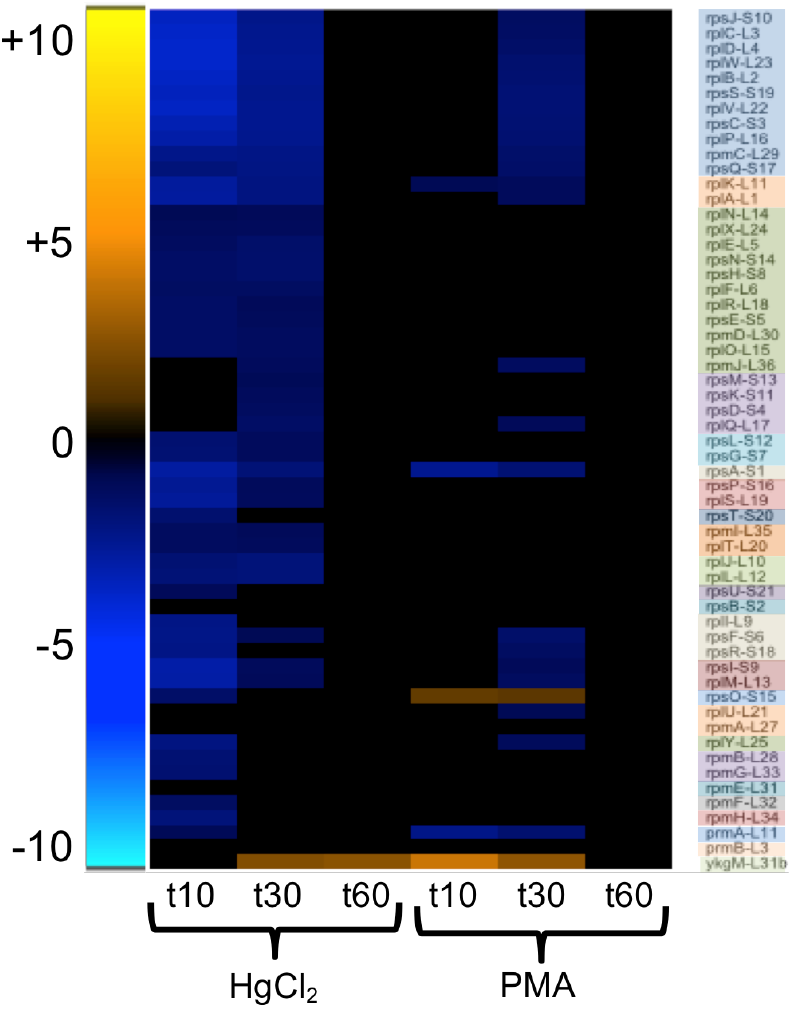
Ribosomal subunit protein genes. Genes are grouped and colored by operon (see larger Figure S11 and Table S13 for details).

Translation initiation and elongation factors were largely unchanged, but expression of all three peptide chain release factor genes were down-regulated for PMA and the ribosome recycling factor (frr) was up-regulated only for PMA, consistent with interruption of translation. Eight tRNA-synthase genes declined with HgCl_2_ but PMA caused only four tRNA-synthase genes to decline and two to increase in expression (Table S14). Both mercurials caused a relative decline in tRNA expression for most amino acids, especially arginine, lysine, methionine, tyrosine, and valine tRNAs. With very few exceptions, ribosome assembly and translation were shut down for up to 30 min by both compounds, but returned to normal levels by 60 min.

### (d) Macromolecular turnover and chaperones

Divalent inorganic mercury can stably crosslink proteins and their subdomains via cysteines, disrupting 3-dimensional structures and allosteric movements [51–53]. Although monovalent PMA cannot cross-link, it forms a bulky adduct with cysteines [23], which may compromise protein folding. The proteases and chaperones of the heat shock response degrade or repair misfolded proteins [54] and we found their expression was increased by both mercurials (Figures 6 and S12 and Table S13). At 10 min, expression of protease genes *lon, clpXP*, and *ftsH* had risen 4-to 6-fold with HgCl_2_ and *lon* and *clpXP*, but not *ftsH*, were up-regulated 3-fold with PMA. HgCl_2_ provoked up-regulation of all 12 heat shock protein (HSP) and chaperone genes by 10 min, but only chaperones *clpB* and *ybbN* mRNAs remained elevated at 30 min. Two other HSP genes, Hsp15 (*hslR*) involved in stalled ribosome recycling and Hsp31 (*hchA*) an amino acid deglycase, were further up-regulated by HgCl_2_ at 60 min. In contrast, at 10 min PMA had up-regulated only five HSPs, increasing to six by 30 min and declining to three by 60 min. The *ibpA* and *ibpB* chaperone genes were among the most highly up-regulated genes for both HgCl_2_ and PMA and persisted throughout recovery.

**Figure 6:**
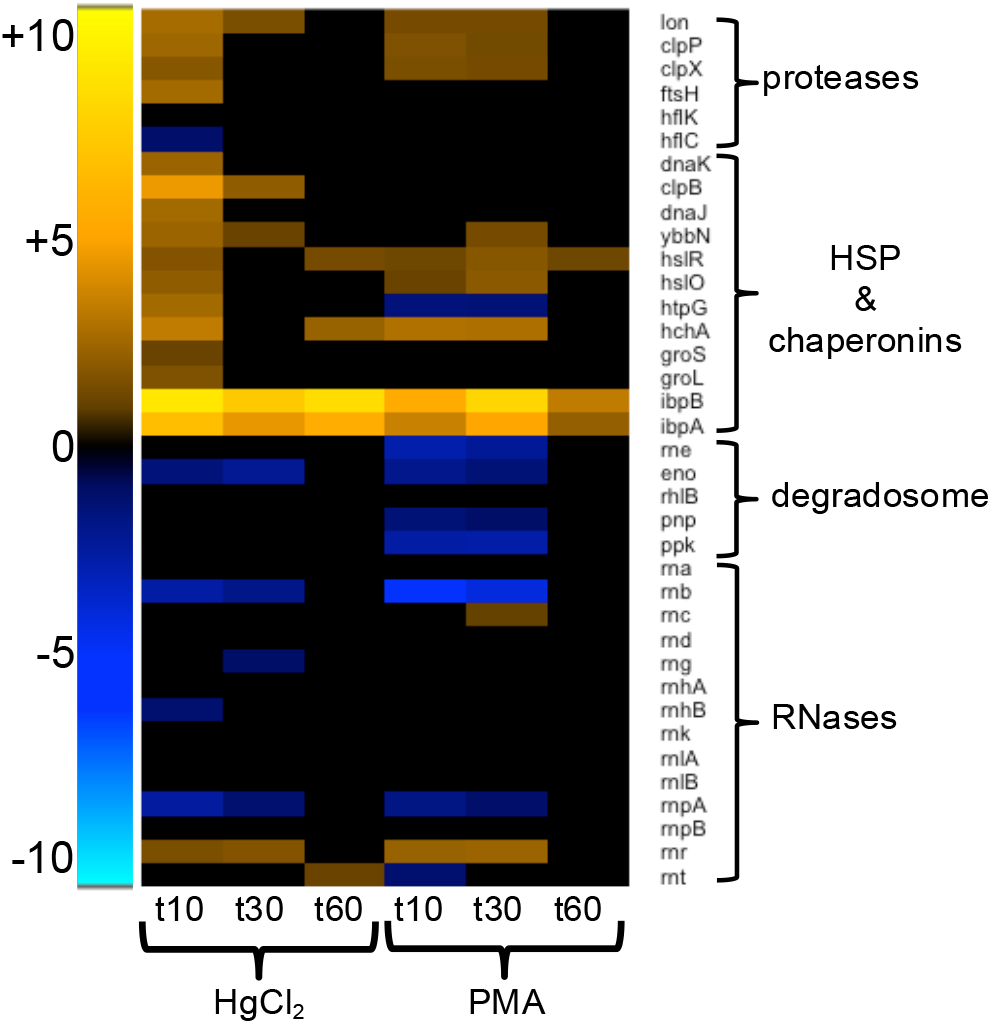
Protein and RNA turnover and repair. (see larger Figure S12 and Table S13 for details).

Of the 16 RNases and RNA processing enzymes only 3 increased: RNase R (3-fold for HgCl_2_ and 5-fold for PMA) for both compounds during first 30 min; RNase III two-fold for PMA at 30 min; and RNase T two-fold for HgCl_2_ at 60 min. The degradasome complex subunit genes (*rne, eno, rhlB, pnp, ppk*) [55] were all down-regulated for PMA during the first 30 min following exposure, except the helicase (*rhlB)*, but only enolase was down-regulated for HgCl_2_ (Figures 6 and S12 and Table S13). Expression of RNase II (*rnb*) was also down-regulated for both compounds, but with relatively greater fold-changes observed for PMA. It is unclear what effect these changes in gene expression could have on RNA turnover and message decay rates while under mercury stress. As with the DNA metabolism genes, expression of the transcriptional apparatus shows that sufficient PMA was taken up to elicit both positive and negative responses distinct from HgCl_2_.

## ii. ENERGY PRODUCTION

### (a) Electron transport chain

Expression of approximately 50% of all electron transport chain (ETC) genes was down-regulated during the first 30 min for HgCl_2_ and PMA, with individual gene responses being very similar for both compounds (Figures 7 and S13 and Table S13).

**Figure 7:**
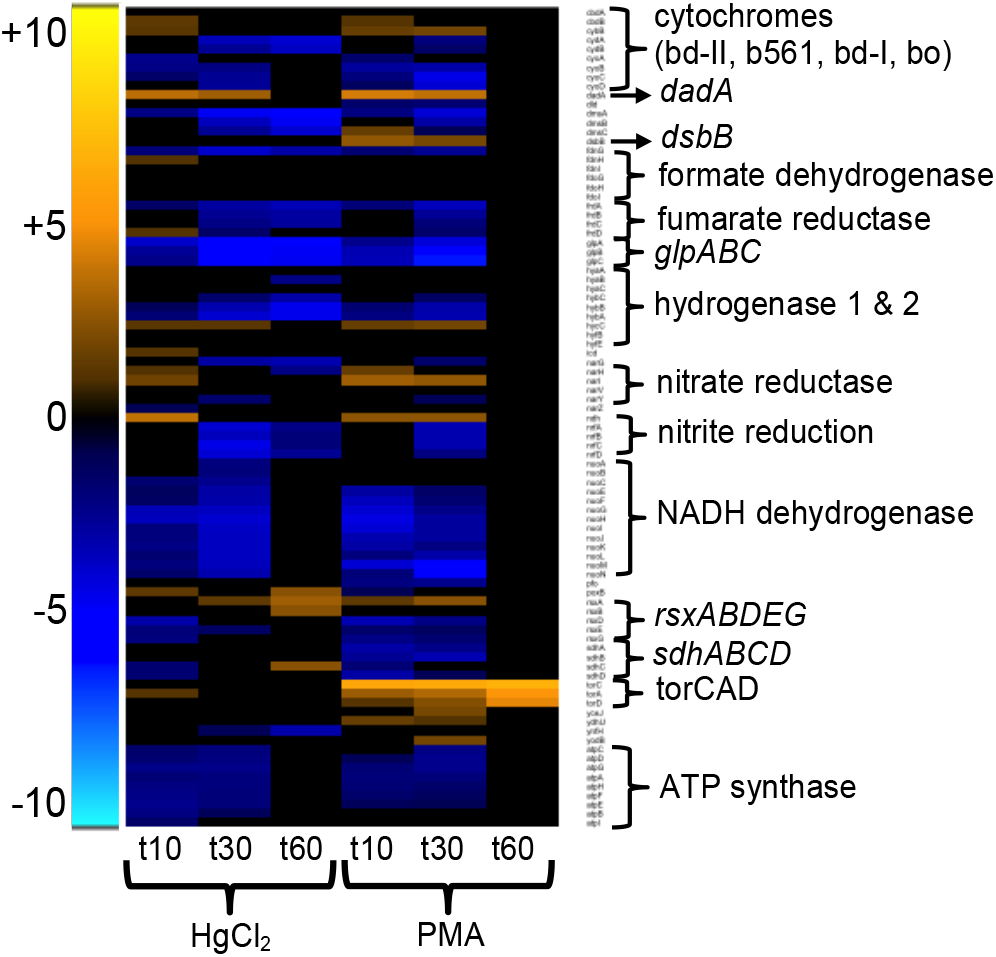
Electron transport chain and ATP-synthase. (see larger Figure S13 and Table S13 for details).

By 60 min, only 26% of these genes were down-regulated for HgCl_2_, and none were down-regulated for PMA. Expression of NADH:unbiquinone oxidoreductase genes was down-regulated by both compounds, with 77% and 100% of these genes being down-regulated at 10 and 30 min, respectively. The ATP-synthase subunit genes were also strongly down-regulated by both mercurials at 10 and 30 min, but normal expression was restored at 60 min.

The *torCAD* locus that encodes the trimethylamine N-oxide anaerobic respiratory system was strongly up-regulated only by PMA exposure. This is likely an artifact of low-level basal expression and the dimethyl sulfoxide used to dissolve PMA. The final DMSO concentration, 0.015% vol/vol (2.1 mM), was not expected to have any biological effect [56] and the anaerobic DMSO reductase genes (*dmsABC*) were down-regulated.

It is unlikely that either the *tor* or *dms* responses affect growth rate [57] or afford protection against either mercurial since over-expressed heme-dependent *torC* may be in the apoprotein form [58].

### (b) Carbon metabolism

Expression of genes for carbon metabolism decreased generally, but there were more up-regulated genes in response to HgCl_2_ and not all steps were affected equally by both mercurials (Figure S14). Expression of five genes of the pentose-phosphate pathway rose in at least one time point for HgCl_2_, but only *pgl* increased for PMA at 10 min. The ribose-5-phosphate isomerase gene (*rpiB*), which is a backup enzyme for the gene product of *rpiA* [59], was up-regulated 40-fold for HgCl_2_ at 10 min; although expression of *rpiA* did not differ from the unexposed cells for either mercurial at any time. Glycolysis responded similarly to both mercurials, with the greatest number of these genes being down-regulated at 30 min. The expression changes in TCA cycle genes were distinct for HgCl_2_ and PMA; six genes were up-regulated in at least one time point for HgCl_2_ and only one was up-regulated for PMA. Expression of several carbohydrate transport genes was down-regulated by both mercurials (Table S6).

### (c) Nicotinamide adenine dinucleotide (NAD)

Expression of genes for nicotinamide adenine dinucleotide (NAD) and NAD-phosphate (NADP) synthesis and turnover pathways was repressed by mercury exposure (Figure S15). The biosynthesis genes were moderately down-regulated, with *nadB* being the only gene down-regulated for both mercurials at all times and *nadA* decreasing for HgCl_2_ at 30 and 60 min and for PMA at 30 min. Expression of the *pncABC* salvage pathway did not change. The NAD reduction pathways were more affected than the NADP reduction pathways, with only *pgi* down-regulated for both mercurials and *edd* down-regulated only for HgCl_2_. The transhydrogenase (*pntAB*) was down-regulated only for PMA at 10 and 30 min. Expression of other genes for NAD to NADH reduction in glycolysis and the TCA cycle were also down-regulated for both mercurials, which reflects the overall decrease in metabolism and energy production pathways.

Globally, redox metabolism declined immediately after exposure and normal gene expression levels were not restored until growth recovered to the pre-exposure rate. KEGG maps created using iPath [60] depict system-wide metabolism changes over time (Figures S16-S18 for HgCl_2_ and Figures S19-S21 for PMA).

## iii. CENTRAL METABOLISM

### (a) Amino acid metabolism and transport

The two mercurials had distinct effects on expression of genes for biosynthesis of amino acids (Figure 8 and S22). Since mercury targets cysteine thiol groups and will deplete the cellular reduced thiol pool, we expected an increase in cysteine and glutathione biosynthesis. Surprisingly, most genes for biosynthesis of these biothiols and for general sulfur metabolism were down-regulated or no different from the unexposed cells, with the exception of up-regulation of *cysE*, which is the first step in the biosynthesis pathway from serine.

**Figure 8:**
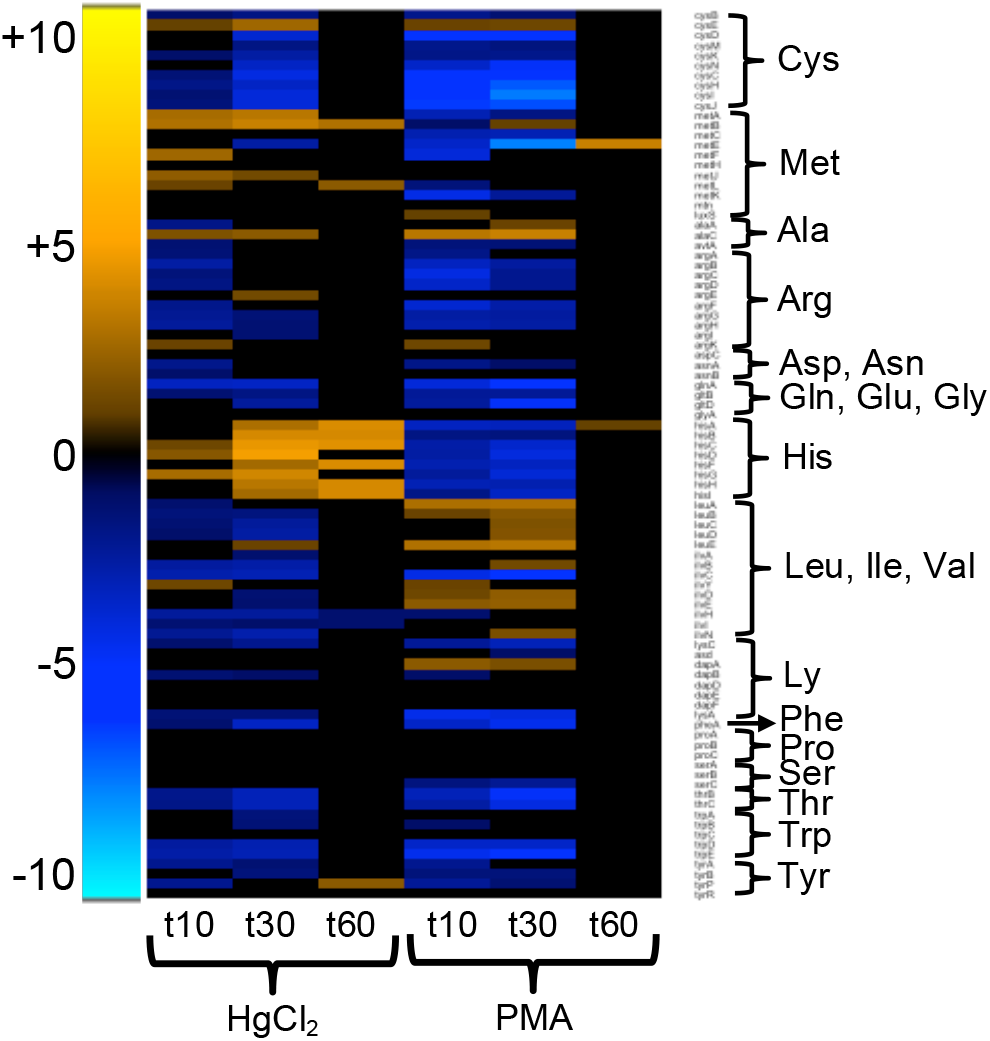
Amino acid biosynthesis. (see larger Figure S22 and Table S13 for details).

Methionine biosynthesis gene expression increased for 7 genes with HgCl_2_, but 11 genes were down-regulated with PMA, especially *metE* dropping 187-fold with PMA at 30 min (Figure 8 and S22). Expression of genes for histidine synthesis also responded differently to each mercurial, rising dramatically with HgCl_2_ at 30 to 60 min.

In contrast, all *his* genes expression dropped with PMA from 10 to 30 min. Genes for the synthesis of leucine, isoleucine, and valine had the opposite response, with most down-regulated with HgCl_2_ but up-regulated with PMA. Expression of other amino acid biosynthetic pathways was largely unchanged or declined with both mercurials. Branched-chain (*livKHMGF)*, dipeptide (*dppABCDF*), and oligopeptide (*oppABCDF*) transporters were also down-regulated for both mercurials, with greater relative negative fold-changes for PMA (Table S5).

### (b) lnorganic ion transport and metallochaperones

Inorganic Hg(II) can displace beneficial thiophilic metals from their native binding sites in proteins, potentially affecting transport and disrupting transition metal homeostasis [23], leading to expression changes for non-ferrous metal cation and oxoanion transporters (Figures 9 and S23 and Table S13), iron homeostasis (Figures 10 and S24 and Table S13) and metal-binding proteins and enzymes (Table S15).

**Figure 9:**
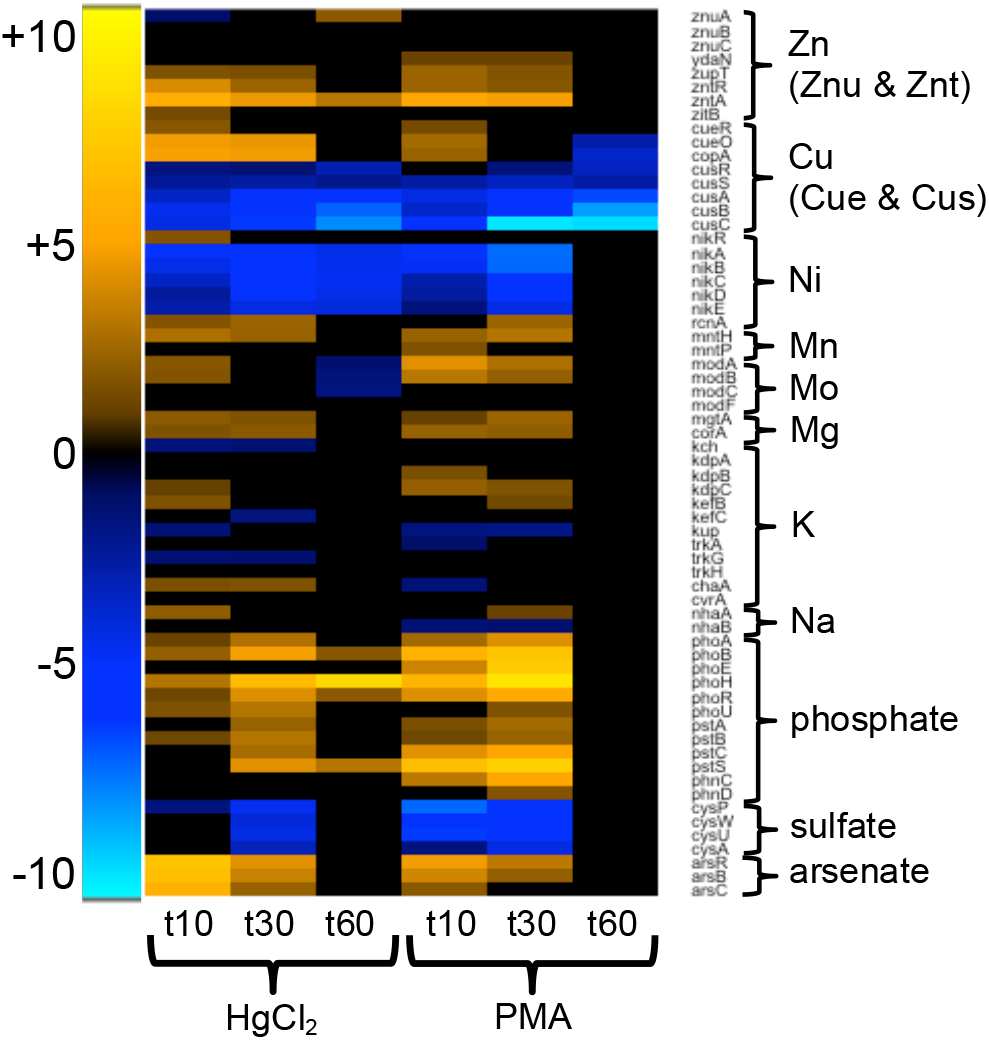
Non-ferrous metals homeostases. (see larger Figure S23 and Table S13 for details).

**Figure 10:**
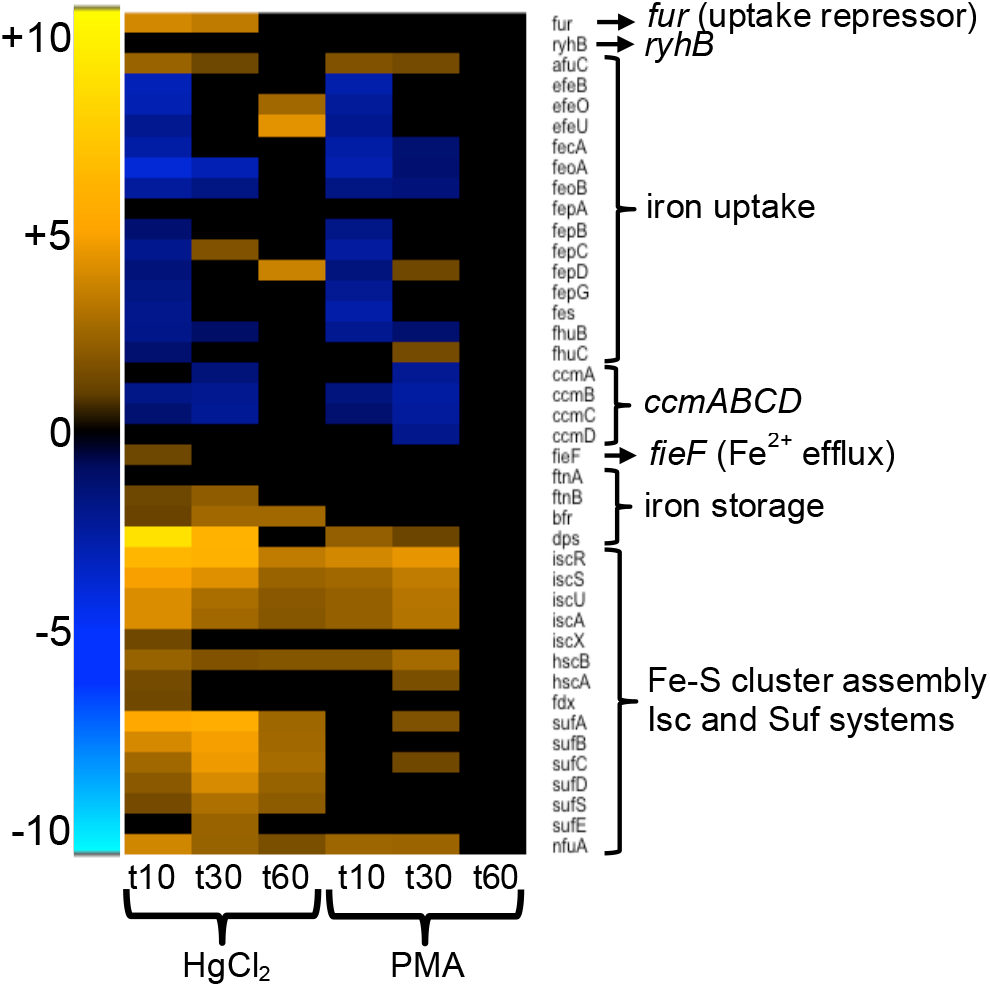
Iron homeostasis. (see larger Figure S24 and Table S13 for details).

Inorganic mercury exposure releases labile iron, which could itself increase oxidative stress via Fenton chemistry under aerobic growth [23, 61]. Most iron uptake pathways declined early for both mercurials, consistent with the observed increase in expression of the Fur repressor. The cytochrome *c* maturation genes that transport heme to the periplasm (*ccmABCDE*) were also down-regulated for both mercurials. The putative ferrous iron and zinc efflux pump, *fieF* [62] increased 2-fold for HgCl_2_ at 10 min only, suggesting it may have a transient role in restoring one or both of these homeostases.

There are two iron-sulfur (Fe-S) cluster assembly pathways in *E. coli* [63, 64]. Expression of the primary Isc system (*iscRSUA*, *hscBA*, and *fdx*) increased strongly for both mercurials, but with greater relative changes for HgCl_2_ (Figures 10 and S24 and Table S13). The secondary Fe-S cluster assembly system *sufABCDSE*, which activates under oxidative stress or iron limiting conditions also increased greatly, but only for HgCl_2_. These transcriptional responses confirm and extend biochemical findings [23] that Fe-S clusters are more vulnerable to inorganic mercury than to organomercurials and the cell quickly tries to repair this damage.

Expression of the zinc transporter *zupT* increased modestly in the 3-to 5-fold range for both mercurials during the first 30 minutes (Figures 9 and S23 and Table S13). In contrast, expression of the P-type ATPase zinc efflux pump, *zntA* [65] increased in the 20-to 40-fold range for both mercurials at 10 and 30 min and the periplasmic Zn-binding protein ZraP was up-regulated throughout recovery. *E. coli* has two copper/silver efflux systems, Cue and Cus [66]. Surprisingly, the Cus system genes (*cusRS*, *cusCFBA*) primarily used under anaerobic conditions were among the most down-regulated genes under PMA exposure. The Cue system consists of the multicopper oxidase, CueO, and a P-type ATPase, CopA, both regulated by the MerR homolog, CueR. Genes *cueO* and *copA* were up-regulated approximately 20-fold with HgCl_2_ at both 10 and 30 min whereas they increased 5-fold with PMA only at 10 min. The nickel uptake system [67] (*nikABCDER*) was also strongly down-regulated under PMA exposure conditions through all times although expression of repressor NikR was unchanged, except for a 3-fold increase with HgCl_2_ at 10 min. Expression of the nickel and cobalt efflux gene, *rcnA*, increased with HgCl_2_ or PMA. Manganese (*mntH)*, and magnesium (*mgtA, corA*) uptake genes increased with both mercurials.

Inorganic anions used by *E. coli* include phosphate, sulfate, and molybdate and the genome also encodes genes for defense against arsenate, which acts as a phosphate mimic (Figures 9 and S23 and Table S13). Expression of the ABC phosphate transport system (*pstSCAB*) genes increased greatly for both mercurials, with PMA-provoked changes up to 165-fold, relative to unexposed condition, for the phosphate binding protein, *pstS*. The two-component phosphate regulatory system, PhoBR, was up-regulated for both mercurials; *phoB* changed up to 22-fold with HgCl_2_ and 105-fold with PMA relative to the unexposed condition. Sulfate and thiosulfate uptake by the ABC transporter (*cysPUWA*) decreased strongly with HgCl_2_ at 30 min and PMA at 10 and 30 min. Expression of molybdate uptake (*modABC*) increased with PMA during the first 30 min but only at 10 min with HgCl_2_. The arsenate resistance operon cannot effect Hg(II) resistance, but was highly induced by both mercurials, perhaps through interacting with the three-cysteine metal-binding site of the ArsR repressor [68].

## iv. SURFACE FUNCTIONS

### (a) Cell wall biogenesis, porins, lps, efflux systems, and electrolyte balance

The transcriptional response of peptidoglycan, membrane biosynthesis, and cell division genes was similar for both mercurials (Figure S25 and Table S13). Expression increased for roughly 20% of lipid biosynthesis genes, including those for cardiolipin, and expression decreased for 20-30% of other lipid-related genes. Transcription of genes for murein synthesis (*murCDEFGIJ*) in particular declined for both mercurials during the first 30 min.

*E. coli* encodes several antibiotic resistance efflux systems that are up-regulated by mercury exposure (Figures 11 and S26 and Table S13). The multiple antibiotic resistance locus (*marRAB*), which increases drug efflux and also limits passive uptake by decreasing porin expression [69], was strongly up-regulated by both mercurials with greater fold-changes observed for PMA, relative to the unexposed condition. Though expression of some porin genes (*ompC*, *ompF*, *ompT*, *ompW*) was repressed, three non-specific porins (*ompG*, *ompL*, *ompN*) were up-regulated only by PMA. Genes from several TolC-dependent antibiotic efflux systems were up-regulated by both mercurials as well, including *acrEF*, *emrD*, *emrKY*, and several *mdt* genes [70]. HgCl_2_ exposure alone also up-regulated two-component sensor genes (*phoQP* at 10 min and *basSR* at 60 min) that regulate genes involved in modification of the cell surface and increase polymyxin resistance [71], but most of these genes were down-regulated or unchanged for PMA.

**Figure 11:**
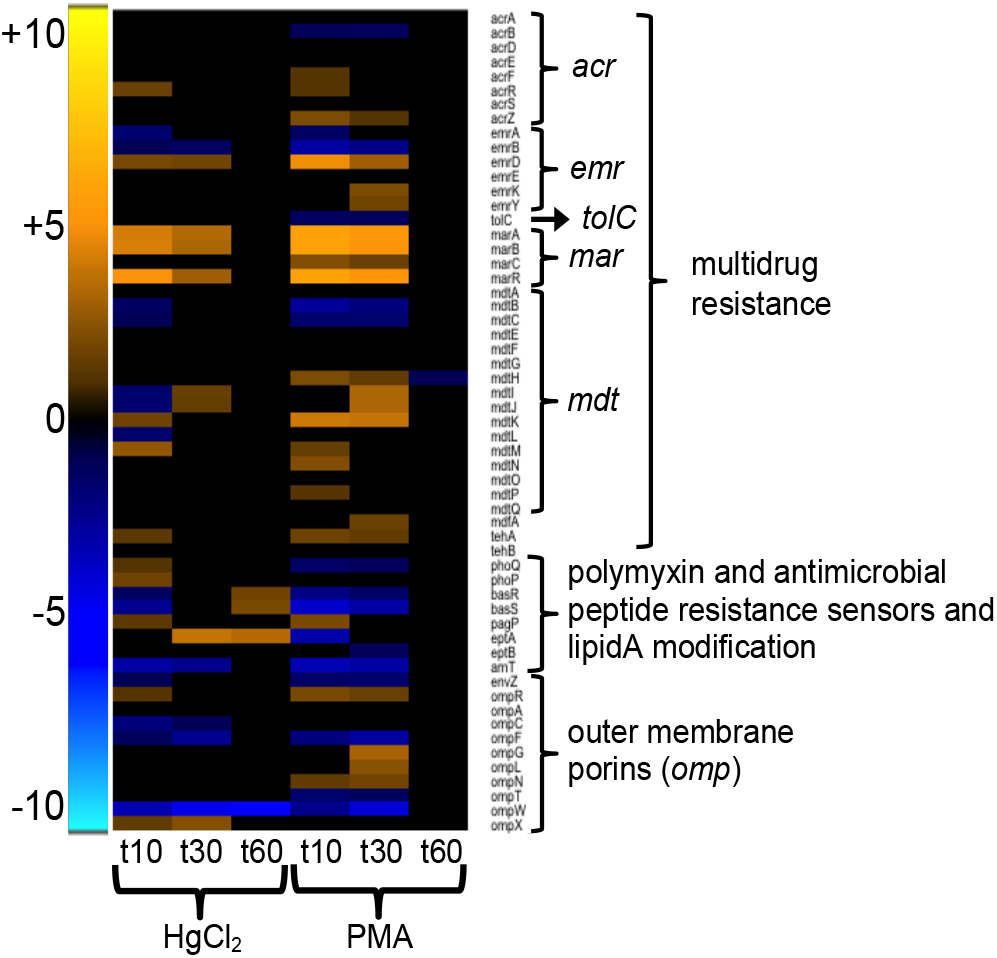
Antibiotic resistance and outer membrane porins. (see larger Figure S26 and Table S13 for details).

The response to osmotic stress and maintenance of electrolyte balance are important membrane functions requiring adaptation in dynamic natural environments. During HgCl_2_ exposure the expression of the sodium antiporter, NhaA, increased 4-fold at 10 min and the calcium/potassium antiporter, ChaA, was up-regulated 3-fold at 10 and 30 min (Figures 9 and S23 and Table S13). In contrast, expression of genes for transport of the major electrolyte, potassium, changed only modestly in some subunits of the *kdp*, *kef*, and *trk* systems, without an obvious response pattern. However, transcription of genes for defense against osmotic stress was uniformly up-regulated; betaine genes (*betABIT and proP), osmBCEFY, and* mechanosensitive channel proteins (*mscL* and *mscS*) increased for both mercurials, as did a putative osmoprotectant ABC permease (yehYXW) [72] only with HgCl_2_ at 30 and 60 min (Table S5).

### (b) Motility and biofilm

Nearly all flagellar component genes were strongly down-regulated for both mercurials, with greater negative fold-changes observed with PMA relative to the unexposed condition (Figures 12 and S27). Only PMA increased expression of fimbriae and curli fiber genes, which alter motility and increase adhesion (Figure 13 and S28) [73]. Fifteen genes up-regulated by PMA exposure were annotated as homologs of FimA, but with unknown function. FimA is the major structural component of fimbriae, but these genes may serve other functions. Motility genes whose expression dropped remained low until 60 min with HgCl_2_, indicating that the structurally and energetically intensive motility systems are very slow to recover.

**Figure 12:**
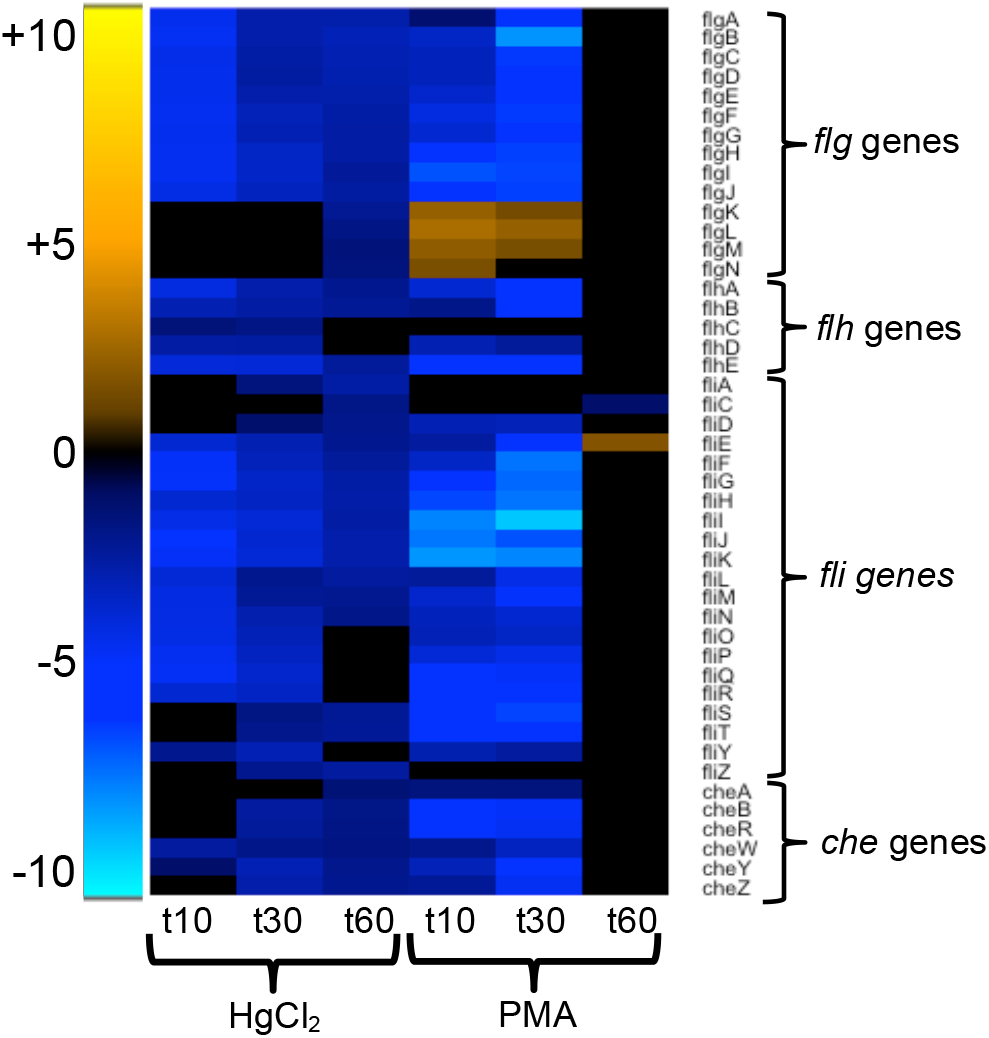
Flagella components and chemotaxis. (see larger Figure S27 and Table S13 for details).

**Figure 13:**
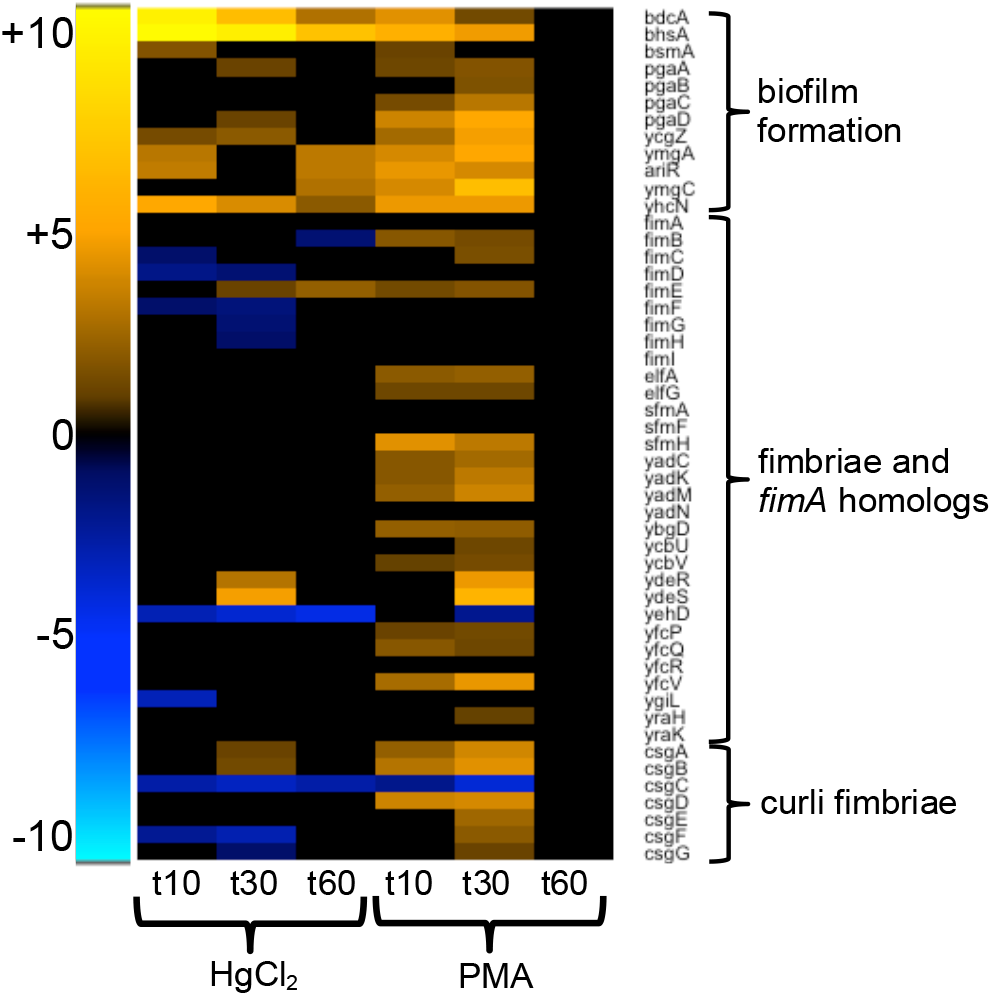
Biofilm formation and fimbriae. (see larger Figure S28 and Table S13 for details).

HgCl_2_ and PMA also provoked expression of several biofilm-related genes (Figures 13 and S28 and Table S13). The *bhsA* and *bdcA* loci were among the most highly up-regulated genes during HgCl_2_ exposure, with a relatively greater fold-change than observed for PMA (Tables 2 and 3). Neither gene is well characterized, but independently each has been found to decrease biofilm formation and increase resistance to external stressors [74, 75]. Only PMA increased expression of genes for poly-β-1,6-N-acetyl-glucosamine (PGA) polysaccharide production [76] and biofilm related genes, *ycgZ*, *ymgA*, *ariA*, *ymgC* [77]. Thus, PMA elicits a broader response that potentially alters the cell surface and may increase adhesion and biofilm formation; in contrast HgCl2 only inhibits motility and does not activate adhesion pathways. It is possible that some changes observed for motility and biofilm related genes following PMA-exposure are an artifact of the DMSO, but other studies suggest that solvent would have no effect or that much higher concentrations than used here would be required to induce these changes [56, 78].

## v. STRESS RESPONSES

### (a) Oxidative stress response and repair

There are two oxidative stress response pathways in *E. coli*, the *oxyRS* and *soxRS* regulons [61, 79]. OxyR, a LysR-family transcriptional regulator, uses a cysteine-pair to sense oxidative damage and regulates 49 genes when oxidized [80]. HgCl_2_ exposure increased expression of 22 OxyR regulon genes at 10 min; these then declined to 13 genes by 60 min (Table S16). In contrast, PMA provoked expression of 16 OxyR regulon genes at 10 and 30 min, but none at 60 min. OxyS, a small non-coding RNA regulated by OxyR, represses *rpoS, fhlACD* and other genes to prevent redundant induction of stress response genes [81]. The *oxyS* gene was among the most highly differentially expressed genes, increasing over 1,000-fold with HgCl_2_ at 10 and 30 min relative to the unexposed condition. Differential expression of *oxyS* was more modest with PMA having a relative increase of 10-fold at 10 min and 6-fold at 30 min.

The SoxRS regulon is the other oxidative stress response system in *E. coli*. SoxR, a MerR-family repressor-activator, uses the oxidation state of 2Fe-2S clusters to respond to superoxide (O_2_^−^) stress and induce transcription of *SoxS* [82–85], which then transcriptionally regulates 53 genes [79, 86] (Table S16). HgCl_2_ or PMA exposure up-regulated 22 or 25 genes, respectively, at 10 min and these had declined to 13 or 0 genes, respectively by 60 min.

Key genes in these oxidative stress regulons differentially expressed upon mercury exposure include the ROS scavengers: catalase (*katG*), alkyl hydroperoxide reductase (*ahpCF*), and superoxide dismutase (*sodA*) (Figures 14 and S29 and Table S13). Thiol homeostasis genes included *gor, grxA*, and *trxC* (Figures 14 and S29 and Table S13). Iron homeostasis and the Fe-S cluster assembly and repair genes (*fur, dps, fldA, fpr, hemH, sufABCDES*, and *yggX*) were also up-regulated. PMA provoked comparatively lower fold-changes than HgCl_2_ for *grxA, trxC, ahpC, dps, fldA, hemH, and yggx*. The manganese uptake protein, *mntH*, plays an important role in ROS resistance [87] and was up-regulated for both mercurials. Oxidation-resistant dehydratase isozymes, *acnA* and *fumC* [88, 89] also increased, but only for HgCl_2_ exposure. Thus, both mercurials triggered the Oxy and Sox oxidative stress responses, but HgCl_2_ elicited greater fold-changes overall than PMA compared to unexposed cells.

**Figure 14:**
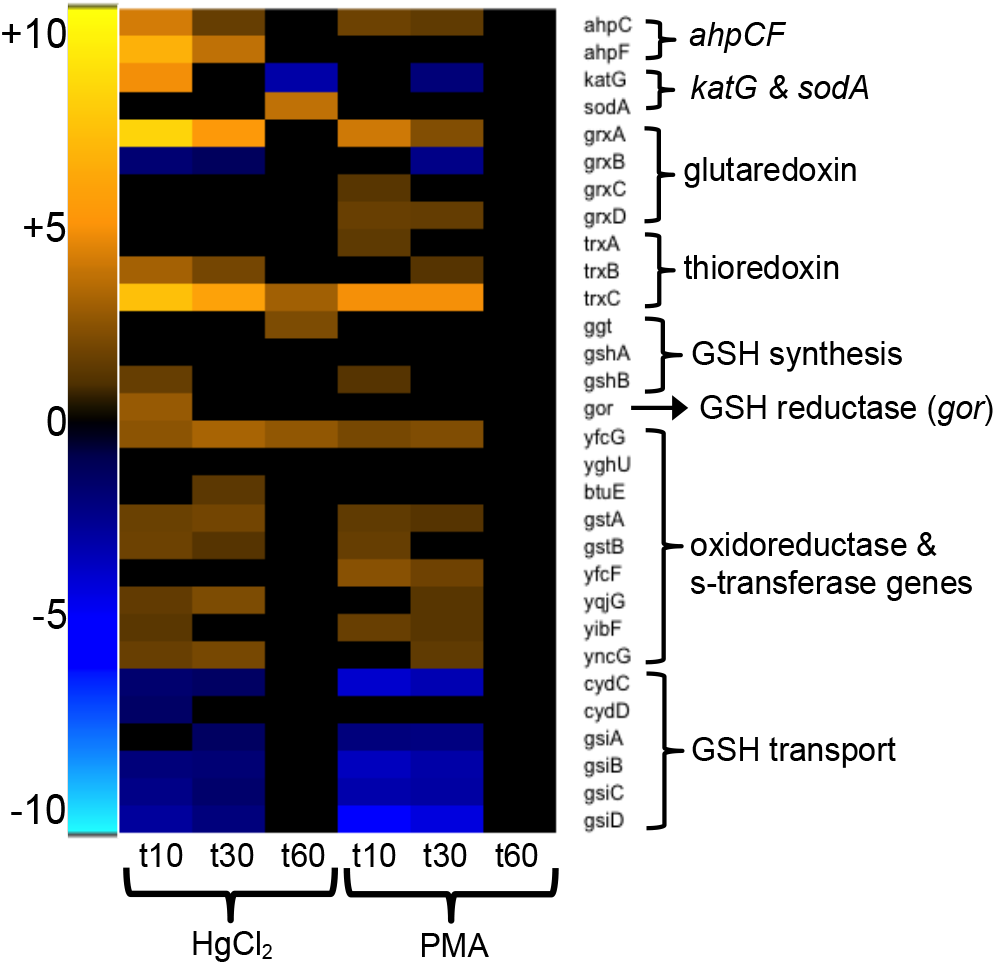
Oxidative stress defense and thiol homeostases. (see larger Figure S29 and Table S13).

Because mercury poisons the cellular thiol pool [23], we expected that regulation of redox homeostasis proteins such as glutaredoxins, thioredoxins and glutathione-related genes would respond to mercury exposure (Figure 14 and S29). Glutaredoxin 1 (*grxA*) expression was up-regulated for both mercurials, with a greater fold-change observed for HgCl_2_ relative to unexposed condition. In contrast, glutaredoxin 2 (*grxB*) was down-regulated for both mercurials, while glutaredoxin 3 (*grxC*) and glutaredoxin 4 (*grxD*) were up-regulated only for PMA. Thioredoxin reductase (*trxB*) was up-regulated 2-fold with PMA only at 30 min, but was up-regulated with HgCl_2_ 8-fold (10 min) and 4fold (30 min). Thioredoxin 1 (*trxA*) expression increased 2-fold only with PMA (10 min); in contrast to thioredoxin 2 (*trxC)*, which was up-regulated for both mercurials, but had greater fold-change relative to the unexposed condition with HgCl_2_ than with PMA. However, the thiol peroxidase (*tpx*) was up-regulated modestly for PMA, but did not change for HgCl_2_. Thus, while each mercurial stresses the cell to maintain redox homeostasis, HgCl_2_ exposure elicited greater responses.

Glutathione (GSH) serves as the cell’s redox buffer and as a scavenger of mercurials (Figures 14 and S29 and Table S13). Surprisingly, expression of GSH biosynthesis and utilization genes increased only modestly. The γ-glutamyltranspeptidase (ggt) increased late (60 min) by 4-fold only with HgCl_2_. GSH synthase (*gshA*) did not change and *gshB* was up-regulated 2- or 3-fold for both mercurials only at 10 min. The GSH importer (*gsiABCD*) may be a salvage pathway to recover GSH and cysteine leaked into the periplasm by CydCD [90, 91], but it was down-regulated by HgCl_2_ or PMA through 30 min. The GSH reductase (*gor*) increased only with HgCl_2_ at 10 min and several GSH S-transferase genes involved in detoxification [92] (*gstA, gstB, yfcF, yqjG, yibF, yncG*) increased with both mercurials. Since all of these proteins have Hg(II)-vulnerable cysteines in their active sites, it is surprising that neither Hg(II) nor PMA-challenged cells provoked increased expression and suggests that their normal mRNA levels are sufficient to replenish them.

### (b) Genes with delayed up-regulation

Genes unchanged at 10 min but differentially expressed at both 30 and 60 min or 60 min alone may be those needed as cells transition out of stasis and towards normal growth (Table S17). For HgCl_2_ exposure, 95 genes were up-regulated and 140 genes were down-regulated that display this delayed response pattern. Approximately half of the up-regulated genes are involved in energy production, transport and metabolism pathways based on COG annotations. Roughly 45% of these delayed HgCl_2_ provoked, up-regulated genes are the same as genes that were differentially expressed during the first 30 min of PMA exposure. This overlap is consistent with slower recovery of growth in HgCl_2_ exposed cells and that some of the same pathways are used for recovery by both compounds. In contrast, for the more quickly recovering PMA exposure, of the genes that showed no change at 10 min only six were up-regulated at 60 min (Table S17). Only two of these delayed-response genes for PMA exposure overlapped with up-regulated genes for HgCl_2_ exposure.

## DISCUSSION

Mercury is a ubiquitous toxicant that serves no biologically beneficial role. Exposure to any form of mercury negatively impacts the health of organisms from microbes to humans. The biological effects of different forms of mercury are often conflated and methylmercury is assumed to be the most toxic form. However, the systemic biochemical and molecular differences between inorganic and organic mercury compounds have yet to be well characterized from exposure through recovery in a single model system.

### BULK DIFFERENTIAL EFFECTS ON GROWTH AND GENE EXPRESSION

The sub-acute mercury exposure conditions used in this study were chosen by identifying a mercury concentration high enough to stop cells from doubling, but low enough to allow restoration of the pre-exposure growth rate within one-hour (~1 generation period) after exposure (Figure 1a). Concentrations below 3 μM of either compound did not consistently inhibit growth and higher concentrations of HgCl_2_ did not allow recovery within the desired time frame. The 3 μM Hg used in this study is well within the range that bacteria can experience chronically from dental amalgam fillings [93] and in highly contaminated environments, such as artisanal gold mining operations [8]. Mercury in tuna is 0.386 ppm compared to the proxy organomercurial, PMA, used here at 3 μM or 0.6 ppm [94].

PMA-exposed cells recovered exponential growth faster (Figure 1) than those with equimolar exposure to HgCl_2_, perhaps owing to lower uptake of PMA (Table S2). However, PMA-exposed cells differentially expressed more genes than HgCl_2_ exposed cells during the first 30 min of exposure (Figure 1b). These results agree with observations in *C. elegans*, where MeHg exposure resulted in four times more DEGs than did HgCl_2_ for all concentrations tested [95]. However, in contrast to *E. coli*, whose growth was inhibited more by inorganic HgCl_2_, in *C. elegans* the effective toxic concentration of methylmercuric chloride was lower than for HgCl_2_ [95]. The large number of significant DEGs in response to mercury exposure is greater than the response to hydrogen-peroxide [96] or nitric-oxide [97], but other types of chemical exposures to organic acids [98] and volatile organic compounds [99] in *E. coli* have resulted in similar numbers of DEGs to mercury exposure and the stress response sigma factor RpoS has been shown to regulate up to 23% of the genome alone [100].

We found that most DEGs peaked at 10 min after exposure for both compounds with HgCl_2_ provoking more down-regulated genes and PMA yielding more up-regulated genes throughout the exposure period (Figure 1b). Even though the optical density (OD) of the HgCl_2_ exposed culture showed no growth recovery from 10 min to 30 min, DEGs decreased by 22%, while PMA-exposed cells over the same period had a moderate increase in OD, but only an 8% decrease in DEGs (Figure 1). In contrast in a eukaryotic system, the livers of HgCl_2_ exposed zebrafish continuously increased in DEGs throughout the observed 96 hour exposure period as mercury accumulated in their cells [101].

In *E. coli*, during the first 30 min post-exposure, 50-70% of both up- and down-regulated genes were the same for both compounds (Figure 2), but at the level of individual genes there were both qualitative and quantitative differences in expression (Table S5), consistent with idiosyncratic transcriptional responses to each compound. The nematode *C. elegans* also manifested distinct and even some opposite transcriptional responses to inorganic and organic mercury exposure in a single end-point microarray experiment [24].

As there are not yet other studies of the global transcriptional response of a bacterium to mercury exposure, on the basis of the findings in eukaryotes and our proteomic work [23] we have organized our observations here into those we had expected and those we did not expect from any yet published work.

### EXPECTED AND UNEXPECTED GENE-SPECIFIC CHANGES

#### Expected transcriptional changes

##### (a) Thiol homeostasis

The millimolar cytosolic pool of glutathione (GSH) can sequester mercurials and thereby protect protein thiols from binding strongly to these soft metal toxicants. However, if the GSH pool becomes depleted by mercury complexation, the cell loses this primary defense mechanism. Since proteins of the stress response and repair pathways all contain active site thiols, often as part of Hg(II)-vulnerable Fe-S centers, it is not obvious how a cell that has lost much of its available thiols to Hg(II) chelation can restore its metabolism. Given this, we expected cysteine and glutathione biosynthesis pathways to be up-regulated. However, cysteine biosynthesis was down-regulated (Figures 8 and S22 and Table S13) and GSH biosynthesis was mostly unchanged (Figures 14 and S29 and Table S13) for both compounds, in contrast to the eukaryotic response to mercury [101–103] and H_2_O_2_ exposure [96], which increase GSH and metallothionein production. However, although thiol biosynthesis did not increase, genes involved in maintaining cellular thiol homeostasis did increase (Figures 14 and S29 and Table S13); thioredoxin (*trxC*) and glutaredoxin (*grxA*) were among the most highly up-regulated genes with HgCl_2_ and PMA exposure. Others have also found in *E. coli* that glutathione reductase increased with HgCl2 and both compounds increased expression of glutathione oxidoreductase and S-transferase genes, which protect against oxidative stress and xenobiotics [104].

##### (b) Iron homeostasis

We also expected inorganic mercury to disrupt iron-sulfur clusters with consequent effects on Fe homeostasis generally [23, 105]. Iron uptake was down-regulated with both mercurials, consistent with excess intracellular free Fe(II) and general oxidative stress, but expression of the uptake repressor (*fur*) was only up-regulated for HgCl_2_ (Figure 10 and S24). *Fur* expression is activated by either OxyR or SoxS [96, 106] and Fur represses Fe uptake pathways with ferrous iron as a co-repressor [107]. Fur can also bind other divalent metals [108], so Hg(II)-Fur might mimic Fe-Fur as an iron uptake repressor under these conditions to limit Fenton-mediated damage from excess iron. Although both mercurials increased expression of the primary Fe-S cluster assembly and repair system (isc), only HgCl_2_ induced the secondary system (*suf*), which is normally induced under oxidative stress or iron limiting conditions [63, 109] (Figures 10 and S24 and Table S13). Also, only HgCl_2_ exposure increased expression of iron storage proteins: ferritin, bacterioferritin, and Dps (Figure 10 and S24). The DNA binding protein Dps which binds free iron to protect DNA from ROS damage [110] was one of the most highly up-regulated genes with HgCl_2_ exposure (Table 1).

##### (c) Oxidative stress response

The known close link between iron homeostasis and oxidative stress [61] explains the large fold-changes observed upon HgCl_2_ exposure in genes that respond to oxidative stress (Table S16) and echoes mercury’s long known stimulation of oxidative damage in rat kidney mitochondria [111]. The small non-coding RNA *oxyS* was the second most highly up-regulated gene upon HgCl_2_ exposure with differential expression more than 100-fold greater than observed for PMA (Table 1). The ROS scavenger *ahpF* was also highly up-regulated, along with *katG* (early) and *sodA* (delayed) but only for HgCl_2_ (Figures 14 and S29 and Table S16). Other ROS resistant enzymes, aconitase A [89] and fumarase C [112] (Table S16) also increased only for HgCl2, as did the manganese-dependent alternative ribonucleotide reductase genes (*nrdHIEF*) [113]. The glutaredoxin-like protein that functions like thioredoxin, *nrdH* [114], was highly up-regulated by HgCl_2_ and might support other thioredoxin and glutaredoxin proteins. These striking differences in gene expression illuminate how *E. coli* modulates expression of specific genes not only to deal with compromised function of specific HgCl_2_ modified proteins, but also to manage the consequent cascade of reactive oxygen species.

##### (d) Heat shock response

Mercurials bound to protein cysteines could disrupt protein folding, subunit assembly, and allosteric movements [52, 115] and inorganic mercury can crosslink neighboring cysteines leading to aggregation [51]. Increased expression of heat shock response genes was expected as a consequence of such anticipated protein misfolding problems [116]. Indeed, expression of heat shock chaperonins and protease genes increased for both mercurials, with more genes up-regulated early in response to HgCl_2_ than to PMA (Figures 6 and S12 and Table S13). Genes for the small chaperone-like proteins, *ibpA* and *ibpB*, were among the most highly increased for both compounds, especially for HgCl_2_ (Table 1), consistent with their role in aiding Lon protease in the degradation of misfolded proteins [117–119].

##### (e) Translational apparatus

Thiophilic Cd^2+^ exposure in *E. coli* has been shown to decrease expression of ribosomal proteins [120]. In our proteomics work (Zink et al. in preparation) we observed fourteen r-proteins (7 for each ribosomal subunit) that formed stable adducts with either PMA or Hg(II), so it was reasonable to expect this to be reflected in transcription of r-proteins. Indeed, HgCl_2_ exposure repressed expression of up to 83% of r-protein genes at 10 min and 74% at 30 min (Figure 5, S11, and Table S14), whereas PMA only transiently repressed expression of 41 % of r-proteins at 30 min. Divalent inorganic mercury’s ability to cross-link proteins may interfere with ribosome assembly resulting in translational feedback and repression of r-proteins transcription. Disruption of ribosome assembly could also contribute to the slower recovery of growth after inorganic Hg(II) exposure.

##### (f) Energy production

The dependence of most energy production pathways on redox-active transition metals and redox-active sulfur compounds made them obvious targets of mercurial disruption, e.g. three ATP-synthase subunits form stable adducts after in vivo exposure to Hg(II) or PMA (Zink et al. in preparation). Expression of genes within this functional category was largely down-regulated early in exposure, with all nine ATPase subunits down-regulated for HgCl_2_ (10 min) and eight down-regulated for PMA (30 min) (Figure 7 and S13). Although others have found that Cd exposure in *E. coli* repressed aerobic energy metabolism genes and induced anaerobic pathways [120], we found that both aerobic and anaerobic energy metabolism were repressed by HgCl_2_ and PMA. Even though expression of the oxygen-sensing *fnr* [121] and *aer* [122, 123] activators of the anaerobic shift were moderately up-regulated for PMA and unchanged for HgCl_2_ (Table S5). Similarly, glucose metabolism genes were also predominantly down-regulated during early periods for both compounds, especially with PMA exposure (Figure S14). Thus, with severely compromised energy production systems, it is not surprising that amino acid, carbohydrate and nucleotide metabolism genes, and the energy-dependent transport of these molecules (Figure 3 and Table S6), are also largely depressed initially.

##### (g) Homeostases of non-ferrous metals

We expected mercurials to disrupt electrolyte balance [23], but expression of the potassium efflux pumps’ subunit genes (*kcpABC, kefBC, and trkAGH*) were not uniformly up-regulated, although the need to restore the pH balance was indicated by transiently increased expression of the H^+^/Na^+^ antiporter (*nhaA*) for both compounds (Figures 9, S23 and Table S5). It may be that normal levels of the proteins involved in maintaining cellular electrolyte balance are sufficient to respond to mercury exposure and a significant change in transcriptional expression is not required for these genes.

Mercury is also expected to disrupt non-ferrous metal homeostasis because it can displace other metals, such as zinc and copper, as enzyme cofactors [124, 125]. Expression of metal uptake genes decreased and of metal efflux genes increased for zinc, copper, nickel and cobalt, with relatively greater fold-changes occurring with HgCl_2_. Zinc efflux by ZntA is regulated by the MerR homolog ZntR, which can respond to Hg(II) [126], but has not been shown to confer resistance to Hg(II) exposure. Up-regulated periplasmic transition metal binding proteins, ZraP [127] and ZinT [128], use histidine residues to coordinate metal binding and all cysteine residues present in either protein are involved in structural disulfide bonds that prohibit availability for binding metals. Thus, these periplasmic metal binding proteins lack requisite thiol ligands [23] to compete for Hg effectively with periplasmic glutathione [90, 129, 130]. Their increased transcription here likely reflects their control by a complex suite of redox and other stress regulators [86] provoked by Hg exposure as reported above. Manganese may protect iron metalloenzymes under oxidative stress conditions [87] and Mn uptake by *mntH*, as part of the OxyR regulon, was correspondingly up-regulated for both compounds in response to mercury-induced oxidative stress.

#### Unexpected transcriptional changes

##### (a) Motility and chemotaxis

Energetically costly flagellar motility and chemotaxis were strongly down-regulated by both mercurials and were among the slowest to recover normal transcription levels (Figures 12 and S27 and Table S13). Motility gene expression is regulated by σ^28^ (*fliA*) and FlhDC [131] and the expression of these two regulatory genes declined with HgCl_2_ but was unchanged with PMA. Repression of motility may occur through sigma factor competition for binding to RNAP between σ^28^ and increased expression of σ^S^ [132] and/or through repression of the flagellar transcriptional activator FlhDC by increased expression of the small ncRNAs *oxyS* and *gadY* [133]. Interestingly, HgCl_2_ exposure impaired locomotion in *C. elegans* [24], although through a very different mechanism of motility from *E. coli*.

##### (b) Surface appendages and biofilm synthesis

Surprisingly, there were large increases in expression of genes involved in biofilm formation and adhesion or dispersal (Figures 13 and S28 and Tables S13). Expression of *bhsA* and *bdcA*, which function in biofilm dispersal or reduced biofilm formation [74, 75], were the first and third, respectively, most up-regulated genes by HgCl_2_ (Table 1) and were also up-regulated for PMA, but with relatively smaller fold-changes (Table 2). Expression of *bhsA* is also up-regulated by other diverse stressors and may decrease cell permeability [75, 134]. PMA exposure especially increased expression of genes for the polysaccharide PGA, which aids in adhesion in biofilm formation [76], and other biofilm formation (*ycgZ*, *ymgA*, *ariA*, *ymgC*) genes [77] (Figures 13 and S28 and Table S13).

Expression of fimbriae (*fim*) and curli fibers (csg), important for adhesion in biofilm formation [73] were also up-regulated only by PMA (Figure 13 and S28), as were 15 of 22 FimA homologs of unknown function. Outer membrane vesicle formation (OMVs) could also play a role in detoxification, since an increase in formation of these vesicles has been associated with heat shock, oxidative stress response, and biofilm formation [135], which are responses up-regulated to varying degrees by both mercurials. These are distinct differences between PMA and HgCl_2_ response. PMA provocation of biofilm formation and adhesion genes might be an artifact of its DMSO solvent, but it is not obvious why HgCl_2_ induces such high increases in biofilm dispersal genes.

##### (c) Phosphate metabolism

Phosphate uptake genes were among the most highly up-regulated genes for PMA exposure (Figure 9, S23, and Table 2). The PhoBR two-component system controls expression of phosphate transport genes, as well as some genes that increase virulence including those for fimbriae and biofilm formation [136]. Since expression of PhoBR and its regulon increases under phosphate limiting conditions [137], it may be that PMA inhibits phosphate uptake by an unknown mechanism, possibly through direct interaction with highly up-regulated PstS or this could be an artifact of DMSO.

##### (d) Amino acid biosynthesis

Expression of most amino acid pathways was down-regulated by both compounds, but a few responded uniquely to each mercurial (Figures 8 and S22 and Table S13). Since methionine auxotrophy occurs under oxidative stress due to ROS susceptibility of methionine synthase (MetE) [138], expression of methionine biosynthesis genes may have increased with HgCl2 due to a stronger oxidative stress response than PMA. However, since PMA did up-regulate some oxidative stress-related genes, it is curious that Met operon expression was down-regulated under PMA exposure. Next to its affinity for cysteine sulfur, Hg(II) binds the imino nitrogen of histidine very strongly [139], so it was intriguing that histidine biosynthesis genes were also up-regulated by HgCl_2_ but down-regulated by PMA. It remains unclear how these differences or the opposite responses for leucine, isoleucine and valine biosynthesis help the cell survive mercurial exposure.

##### (e) Miscellaneous genes

Multiple antibiotic efflux systems and polymyxin resistance surface modifications were up-regulated by HgCl_2_ exposure, and even more so by PMA (Figures 11 and S26 and Table S13). Chronic mercury exposure contributes to the spread of multiple antibiotic resistant bacteria through co-selection of plasmid-borne antibiotic and mercury resistance genes [140, 141]. Increased expression of antibiotic resistance and surface components hint that low-level mercury exposure could prime cells for increased antibiotic resistance. However, the ubiquity of plasmid- and transposon-borne Hg resistance loci suggests that expression of these chromosomal genes offers insufficient protection against the antibiotic or mercurial levels encountered in clinical practice.

A handful of vestigial e14 or CPS-53 prophage genes were up-regulated by HgCl_2_ (Table 1) or PMA (Table S5), respectively. Some are known to increase resistance to osmotic, oxidative, and acid stressors [142, 143], but their roles and mechanisms have not been well defined.

##### (f) Differential expression of genes required for the same functional protein complex

In many instances we observed that transcripts for subunits of the same enzyme, protein complex, or component of a tightly articulated pathway were differentially expressed. In some cases these proteins lie in distinct transcripts, which may experience different turnover rates and in other cases the differences could be due to transcriptional polarity. We have chosen not to deal explicitly with such paradoxes in this work, which is sufficiently complex as it is, but will address them in future work.

## CONCLUSIONS

The effects of mercury exposure in multicellular organisms have long been studied at the physiological level but a global, fine grained understanding of the differences in the precise biochemical sequelae of inorganic and organic mercury exposure has been lacking. This study is the first to examine not only the global transcriptional response differences between inorganic mercury (HgCl_2_) and an organomercurial (phenylmercuric acetate) in a model microorganism, but also first to examine longitudinally how the cell recovers from these chemically distinct compounds. Taken together with global identification of vulnerable protein targets (Zink et al. in preparation) and of damage to thiol and metal ion homeostasis upon acute mercurial exposure [23], the current work provides a quantitative systems-level description of the effects of *in vivo* mercury exposure in *E. coli*. What was striking and most challenging with this study was the breadth and diversity of the systems whose expression was affected by these two chemically distinct mercurials. Sub-acute exposure influenced expression of ~45% of all genes with many distinct responses for each compound, reflecting differential biochemical damage by each mercurial and the corresponding resources available for repair. Energy production, intermediary metabolism and most uptake pathways were initially down-regulated by both mercurials, but nearly all stress response systems were up-regulated early by at least one compound. These results echo the wide functional variety of proteins stably modified by these mercurials owing to the widespread occurrence of cysteines found in nearly all *E. coli* proteins. Microbiome studies are rapidly unveiling the importance of commensal bacteria to the health of all higher organisms. Our findings in this model commensal organism provide insights into how chronic mercury exposure might affect such complex microbial communities and, consequently, the health of the host. This work also serves as a foundation for studies now underway of how the widely found mobile Hg resistance (*mer*) locus assists the cell in recovery from Hg exposure.

## LIST OF ABBREVIATIONS

Hg: = mercury
HgCl_2_: = mercuric chloride
PMA: = phenylmercuric acetate
PhHg: = phenylmercury
DMSO: = dimethyl sulfoxide
LB: = Luria-Bertani medium
NM3: = Neidhardt MOPS minimal medium
NGS: = next generation sequencing
DEGs: = differentially expressed genes
CVAA: = cold vapor atomic absorption
GSH: = glutathione
Cys: = cysteine
MDR: = multidrug resistance
ncRNA: = non-coding RNA
COGs: = clusters of orthologous groups
GOFs: = gene-ontology functions
RNAP: = RNA polymerase
HSP: = heat shock protein
ETC: = electron transport chain
PGA: = poly-β-1,6-N-acetyl-glucosamine

## DECLARATIONS

### Ethics approval and consent to participate

Not applicable

### Consent for publication

Not applicable

### Availability of data and materials

The tabulated datasets supporting the conclusions of this article are included as additional files. The read counts and raw sequence data (.fastq) are stored and available to the public through the Gene Expression Omnibus database (http://www.ncbi.nlm.nih.gov/geo/) with accession ID: GSE95575.

### Authors’ contributions

SL conceived and designed experiments, prepared biological samples, extracted ribosomal depleted RNA for RNA-Seq, performed all data analysis, and drafted manuscript. AOS was a major contributor in experimental design, feedback on data analysis, and in editing the manuscript. All authors read and approved the manuscript.

## ACKNOWLEDGMENTS

We thank Roger Nilsen at the Georgia Genomics Facility for library preparation and Sharron Crane at Rutgers University for Hg content analysis. We also thank Bryndan Durham, Brandon Satinsky, Mary Ann Moran, Michael K. Johnson, Harry Dailey, Timothy Hoover, Anna Karls, Alexander Johs, Jerry Parks, Susan Miller, and Andrew Wiggins who have provided feedback and discussion on this work. This work was supported by US Department of Energy awards ER64408 and ER65286 to AOS.

## REFERENCES

1. Driscoll CT, Mason RP, Chan HM, Jacob DJ, Pirrone N. Mercury as a global pollutant: sources, pathways, and effects. Environ Sci Technol. 2013;47(10):4967–4983.

2. Pirrone N, Cinnirella S, Feng X, Finkelman RB, Friedli HR, Leaner J, Mason R, Mukherjee AB, Stracher G, Streets DG et al: Global Mercury Emissions to the Atmosphere from Natural and Anthropogenic Sources. In: Mason R, Pirrone N, editors. Mercury Fate and Transport in the Global Atmosphere: Emissions, Measurements and Models. Boston, MA: Springer US;2009. p. 1–47.

3. Mason RP, Fitzgerald WF, Morel FMM. The Biogeochemical Cycling of Elemental Mercury - Anthropogenic Influences. Geochimica Et Cosmochimica Acta. 1994;58:3191–3198.

4. Barkay T, Miller SM, Summers AO. Bacterial mercury resistance from atoms to ecosystems. FEMS Microbiol Rev. 2003;27(2–3):355–384.

5. Crinnion WJ. Environmental medicine, part three: long-term effects of chronic low-dose mercury exposure. Altern Med Rev. 2000;5(3):209–223.

6. Richardson GM, Wilson R, Allard D, Purtill C, Douma S, Graviere J. Mercury exposure and risks from dental amalgam in the US population, post-2000. The Science of the total environment. 2011;409(20):4257–4268.

7. Diez S. Human health effects of methylmercury exposure. Rev Environ Contam Toxicol. 2009;198:111–132.

8. Malm O. Gold mining as a source of mercury exposure in the Brazilian Amazon. Environ Res. 1998;77(2):73–78.

9. Clarkson TW, Magos L. The toxicology of mercury and its chemical compounds. Crit Rev Toxicol. 2006;36(8):609–662.

10. Hintelmann H, Hempel M, Wilken RD. Observation of unusual organic mercury species in soils and sediments of industrially contaminated sites. Environ Sci Technol. 1995;29(7):1845–1850.

11. Tchounwou PB, Ayensu WK, Ninashvili N, Sutton D. Environmental exposure to mercury and its toxicopathologic implications for public health. Environ Toxicol. 2003; 18(3): 149–175.

12. Davidson PW, Myers GJ, Weiss B. Mercury exposure and child development outcomes. Pediatrics. 2004;113(Suppl 4):1023–1029.

13. Valko M, Morris H, Cronin MT. Metals, toxicity and oxidative stress. Curr Med Chem. 2005; 12(10):1161–1208.

14. Yorifuji T, Tsuda T, Takao S, Harada M. Long-term exposure to methylmercury and neurologic signs in Minamata and neighboring communities. Epidemiology. 2008;19(1):3–9.

15. Zahir F, Rizwi SJ, Haq SK, Khan RH. Low dose mercury toxicity and human health. Environ Toxicol Pharmacol. 2005;20(2):351–360.

16. Cheesman BV, Arnold AP, Rabenstein DL. Nuclear magnetic resonance studies of the solution chemistry of metal complexes. 25. Hg(thiol)3 complexes and HG(II)-thiol ligand exchange kinetics. J Am Chem Soc. 1988;110:6359–6364.

17. Oram PD, Fang X, Fernando Q, Letkeman P, Letkeman D. The formation of constants of mercury(II)--glutathione complexes. Chemical research in toxicology. 1996;9(4):709–712.

18. Labunskyy VM, Hatfield DL, Gladyshev VN. Selenoproteins: molecular pathways and physiological roles. Physiol Rev. 2014;94(3):739–777.

19. Pal PB, Pal S, Das J, Sil PC. Modulation of mercury-induced mitochondria-dependent apoptosis by glycine in hepatocytes. Amino Acids. 2012;42(5):1669–1683.

20. Shenker BJ, Guo TL, Shapiro IM. Mercury-induced apoptosis in human lymphoid cells: evidence that the apoptotic pathway is mercurial species dependent. Environ Res. 2000;84(2):89–99.

21. Jomova K, Valko M. Advances in metal-induced oxidative stress and human disease. Toxicology. 2011;283(2–3):65–87.

22. Polacco BJ, Purvine SO, Zink EM, Lavoie SP, Lipton MS, Summers AO, Miller SM. Discovering mercury protein modifications in whole proteomes using natural isotope distributions observed in liquid chromatography-tandem mass spectrometry. Mol Cell Proteomics. 2011; 10(8):M110.004853.

23. LaVoie SP, Mapolelo DT, Cowart DM, Polacco BJ, Johnson MK, Scott RA, Miller SM, Summers AO. Organic and inorganic mercurials have distinct effects on cellular thiols, metal homeostasis, and Fe-binding proteins in Escherichia coli. J Biol Inorg Chem. 2015;20(8):1239–1251.

24. McElwee MK, Ho LA, Chou JW, Smith MV, Freedman JH. Comparative toxicogenomic responses of mercuric and methyl-mercury. BMC Genomics. 2013;14:698.

25. Neidhardt FC, Bloch PL, Smith DF. Culture medium for enterobacteria. J Bacteriol. 1974; 119(3):736–747.

26. Stead MB, Agrawal A, Bowden KE, Nasir R, Mohanty BK, Meagher RB, Kushner SR. RNAsnap: a rapid, quantitative and inexpensive, method for isolating total RNA from bacteria. Nucleic acids research. 2012;40(20):e156.

27. Langmead B, Salzberg SL. Fast gapped-read alignment with Bowtie 2. Nature methods. 2012;9(4):357–359.

28. Li H, Handsaker B, Wysoker A, Fennell T, Ruan J, Homer N, Marth G, Abecasis G, Durbin R, Genome Project Data Processing S. The Sequence Alignment/Map format and SAMtools. Bioinformatics. 2009;25(16):2078–2079.

29. Anders S, Pyl PT, Huber W. HTSeq--a Python framework to work with high-throughput sequencing data. Bioinformatics. 2015;31(2):166–169.

30. Hardcastle TJ, Kelly KA. baySeq: empirical Bayesian methods for identifying differential expression in sequence count data. BMC bioinformatics. 2010;11:422.

31. Hamlett NV, Landale EC, Davis BH, Summers AO. Roles of the Tn21 merT, merP, and merC gene products in mercury resistance and mercury binding. J Bacteriol. 1992; 174(20):6377–6385.

32. Summers AO, Lewis E. Volatilization of mercuric chloride by mercury-resistant plasmid-bearing strains of Escherichia coli, Staphylococcus aureus, and Pseudomonas aeruginosa. J Bacteriol. 1973;113(2):1070–1072.

33. Summers AO, Silver S. Mercury resistance in a plasmid-bearing strain of Escherichia coli. J Bacteriol. 1972;112(3):1228–1236.

34. Schaechter M, Santomassino KA. Lysis of Escherichia coli by sulfhydryl-binding reagents. J Bacteriol. 1962;84:318–325.

35. Janssen BD, Hayes CS. The tmRNA ribosome-rescue system. Adv Protein Chem Struct Biol. 2012;86:151–191.

36. Legendre P. Ward’s Hierarchical Agglomerative Clustering Method: Which Algorithms Implement Ward’s Criterion? Journal of Classification. 2014;31(3):274–295.

37. Szklarczyk D, Franceschini A, Wyder S, Forslund K, Heller D, Huerta-Cepas J, Simonovic M, Roth A, Santos A, Tsafou KP et al. STRING v10: protein-protein interaction networks, integrated over the tree of life. Nucleic acids research. 2015;43(Database issue):D447–452.

38. von Mering C, Jensen LJ, Snel B, Hooper SD, Krupp M, Foglierini M, Jouffre N, Huynen MA, Bork P. STRING: known and predicted protein-protein associations, integrated and transferred across organisms. Nucleic acids research. 2005;33(Database issue):D433–437.

39. Arunasri K, Adil M, Khan PA, Shivaji S. Global gene expression analysis of longterm stationary phase effects in E. coli K12 MG1655. PLoS One. 2014;9(5):e96701.

40. Chang DE, Smalley DJ, Conway T. Gene expression profiling of Escherichia coli growth transitions: an expanded stringent response model. Mol Microbiol. 2002;45(2):289–306.

41. Kuzminov A. Recombinational repair of DNA damage in Escherichia coli and bacteriophage lambda. Microbiol Mol Biol Rev. 1999;63(4):751–813.

42. Lusetti SL, Cox MM. The bacterial RecA protein and the recombinational DNA repair of stalled replication forks. Annu Rev Biochem. 2002;71:71–100.

43. Salgado H, Martinez-Flores I, Lopez-Fuentes A, Garcia-Sotelo JS, Porron-Sotelo L, Solano H, Muniz-Rascado L, Collado-Vides J. Extracting regulatory networks of Escherichia coli from RegulonDB. Methods Mol Biol. 2012;804:179–195.

44. Salgado H, Peralta-Gil M, Gama-Castro S, Santos-Zavaleta A, Muniz-Rascado L, Garcia-Sotelo JS, Weiss V, Solano-Lira H, Martinez-Flores I, Medina-Rivera A et al. RegulonDB v8.0: omics data sets, evolutionary conservation, regulatory phrases, cross-validated gold standards and more. Nucleic acids research. 2013;41 (Database issue): D203-213.

45. Tramonti A, Visca P, De Canio M, Falconi M, De Biase D. Functional characterization and regulation of gadX, a gene encoding an AraC/XylS-like transcriptional activator of the Escherichia coli glutamic acid decarboxylase system. J Bacteriol. 2002;184(10):2603–2613.

46. Nishino K, Senda Y, Yamaguchi A. The AraC-family regulator GadX enhances multidrug resistance in Escherichia coli by activating expression of mdtEF multidrug efflux genes. J Infect Chemother. 2008;14(1):23–29.

47. Tucker DL, Tucker N, Ma Z, Foster JW, Miranda RL, Cohen PS, Conway T. Genes of the GadX-GadW regulon in Escherichia coli. J Bacteriol. 2003;185(10):3190–3201.

48. Lemke JJ, Sanchez-Vazquez P, Burgos HL, Hedberg G, Ross W, Gourse RL. Direct regulation of Escherichia coli ribosomal protein promoters by the transcription factors ppGpp and DksA. Proc Natl Acad Sci U S A. 2011;108(14):5712–5717.

49. Nomura M, Gourse R, Baughman G. Regulation of the synthesis of ribosomes and ribosomal components. Annu Rev Biochem. 1984;53:75–117.

50. Wilson DN, Nierhaus KH. The weird and wonderful world of bacterial ribosome regulation. Crit Rev Biochem Mol Biol. 2007;42(3):187–219.

51. Soskine M, Steiner-Mordoch S, Schuldiner S. Crosslinking of membrane-embedded cysteines reveals contact points in the EmrE oligomer. Proc Natl Acad Sci U S A. 2002;99(19): 12043–12048.

52. Imesch E, Moosmayer M, Anner BM. Mercury weakens membrane anchoring of Na-K-ATPase. The American journal of physiology. 1992;262(5):F837–842.

53. Carvalho CM, Chew EH, Hashemy SI, Lu J, Holmgren A. Inhibition of the human thioredoxin system. A molecular mechanism of mercury toxicity. The Journal of biological chemistry. 2008;283(18): 11913–11923.

54. Georgopoulos C, Welch WJ. Role of the major heat shock proteins as molecular chaperones. Annu Rev Cell Biol. 1993;9:601–634.

55. Carpousis AJ. The RNA degradosome of Escherichia coli: an mRNA-degrading machine assembled on RNase E. Annu Rev Microbiol. 2007;61:71–87.

56. Lim JY, May JM, Cegelski L. Dimethyl sulfoxide and ethanol elicit increased amyloid biogenesis and amyloid-integrated biofilm formation in Escherichia coli. Appl Environ Microbiol. 2012;78(9):3369–3378.

57. Markarian SA, Poladyan AA, Kirakosyan GR, Trchounian AA, Bagramyan KA. Effect of diethylsulphoxide on growth, survival and ion exchange of Escherichia coli. Lett Appl Microbiol. 2002;34(6):417–421.

58. Ansaldi M, Bordi C, Lepelletier M, Mejean V. TorC apocytochrome negatively autoregulates the trimethylamine N-oxide (TMAO) reductase operon in Escherichia coli. Mol Microbiol. 1999;33(2):284–295.

59. Sorensen KI, Hove-Jensen B. Ribose catabolism of Escherichia coli: characterization of the rpiB gene encoding ribose phosphate isomerase B and of the rpiR gene, which is involved in regulation of rpiB expression. J Bacteriol. 1996; 178(4): 1003–1011.

60. Yamada T, Letunic I, Okuda S, Kanehisa M, Bork P. iPath2.0: interactive pathway explorer. Nucleic acids research. 2011;39(Web Server issue):W412-415.

61. Imlay JA. The molecular mechanisms and physiological consequences of oxidative stress: lessons from a model bacterium. Nat Rev Microbiol. 2013;11(7):443–454.

62. Grass G, Otto M, Fricke B, Haney CJ, Rensing C, Nies DH, Munkelt D. FieF (YiiP) from Escherichia coli mediates decreased cellular accumulation of iron and relieves iron stress. Arch Microbiol. 2005;183(1):9–18.

63. Bandyopadhyay S, Chandramouli K, Johnson MK. Iron-sulfur cluster biosynthesis. Biochem Soc Trans. 2008;36(Pt 6):1112–1119.

64. Johnson DC, Dean DR, Smith AD, Johnson MK. Structure, function, and formation of biological iron-sulfur clusters. Annu Rev Biochem. 2005;74:247–281.

65. Rensing C, Mitra B, Rosen BP. The zntA gene of Escherichia coli encodes a Zn(II)-translocating P-type ATPase. Proc Natl Acad Sci U S A. 1997;94(26):14326–14331.

66. Outten FW, Huffman DL, Hale JA, O’Halloran TV. The independent cue and cus systems confer copper tolerance during aerobic and anaerobic growth in Escherichia coli. The Journal of biological chemistry. 2001;276(33):30670–30677.

67. Eitinger T, Mandrand-Berthelot MA. Nickel transport systems in microorganisms. Arch Microbiol. 2000;173(1):1–9.

68. Qin J, Fu HL, Ye J, Bencze KZ, Stemmler TL, Rawlings DE, Rosen BP. Convergent evolution of a new arsenic binding site in the ArsR/SmtB family of metalloregulators. The Journal of biological chemistry. 2007;282(47):34346–34355.

69. Alekshun MN, Levy SB. The mar regulon: multiple resistance to antibiotics and other toxic chemicals. Trends Microbiol. 1999;7(10):410–413.

70. Nishino K, Yamada J, Hirakawa H, Hirata T, Yamaguchi A. Roles of TolC-dependent multidrug transporters of Escherichia coli in resistance to beta-lactams. Antimicrob Agents Chemother. 2003;47(9):3030–3033.

71. Raetz CR, Reynolds CM, Trent MS, Bishop RE. Lipid A modification systems in gram-negative bacteria. Annu Rev Biochem. 2007;76:295–329.

72. Checroun C, Gutierrez C. Sigma(s)-dependent regulation of yehZYXW, which encodes a putative osmoprotectant ABC transporter of Escherichia coli. FEMS Microbiol Lett. 2004;236(2):221–226.

73. Van Houdt R, Michiels CW. Role of bacterial cell surface structures in Escherichia coli biofilm formation. Res Microbiol. 2005;156(5–6):626–633.

74. Ma Q, Zhang G, Wood TK. Escherichia coli BdcA controls biofilm dispersal in Pseudomonas aeruginosa and Rhizobium meliloti. BMC Res Notes. 2011;4:447.

75. Zhang XS, Garcia-Contreras R, Wood TK. YcfR (BhsA) influences Escherichia coli biofilm formation through stress response and surface hydrophobicity. J Bacteriol. 2007; 189(8):3051–3062.

76. Itoh Y, Rice JD, Goller C, Pannuri A, Taylor J, Meisner J, Beveridge TJ, Preston JF, 3rd, Romeo T. Roles of pgaABCD genes in synthesis, modification, and export of the Escherichia coli biofilm adhesin poly-beta-1,6-N-acetyl-D-glucosamine. J Bacteriol. 2008; 190(10):3670–3680.

77. Lee J, Page R, Garcia-Contreras R, Palermino JM, Zhang XS, Doshi O, Wood TK, Peti W. Structure and function of the Escherichia coli protein YmgB: a protein critical for biofilm formation and acid-resistance. J Mol Biol. 2007;373(1):11–26.

78. Cegelski L, Pinkner JS, Hammer ND, Cusumano CK, Hung CS, Chorell E, Aberg V, Walker JN, Seed PC, Almqvist F et al. Small-molecule inhibitors target Escherichia coli amyloid biogenesis and biofilm formation. Nat Chem Biol. 2009;5(12):913–919.

79. Seo SW, Kim D, Szubin R, Palsson BO. Genome-wide Reconstruction of OxyR and SoxRS Transcriptional Regulatory Networks under Oxidative Stress in Escherichia coli K-12 MG1655. Cell Rep. 2015;12(8):1289–1299.

80. Choi H, Kim S, Mukhopadhyay P, Cho S, Woo J, Storz G, Ryu SE. Structural basis of the redox switch in the OxyR transcription factor. Cell. 2001; 105(1):103–113.

81. Altuvia S, Zhang A, Argaman L, Tiwari A, Storz G. The Escherichia coli OxyS regulatory RNA represses fhlA translation by blocking ribosome binding. EMBO J. 1998; 17(20):6069–6075.

82. Amabile-Cuevas CF, Demple B. Molecular characterization of the soxRS genes of Escherichia coli: two genes control a superoxide stress regulon. Nucleic acids research. 1991;19(16):4479–4484.

83. Greenberg JT, Monach P, Chou JH, Josephy PD, Demple B. Positive control of a global antioxidant defense regulon activated by superoxide-generating agents in Escherichia coli. Proc Natl Acad Sci U S A. 1990;87(16):6181–6185.

84. Tsaneva IR, Weiss B. soxR, a locus governing a superoxide response regulon in Escherichia coli K-12. J Bacteriol. 1990;172(8):4197–4205.

85. Watanabe S, Kita A, Kobayashi K, Miki K. Crystal structure of the [2Fe-2S] oxidative-stress sensor SoxR bound to DNA. Proc Natl Acad Sci U S A. 2008;105(11):4121–4126.

86. Keseler IM, Mackie A, Peralta-Gil M, Santos-Zavaleta A, Gama-Castro S, Bonavides-Martinez C, Fulcher C, Huerta AM, Kothari A, Krummenacker M et al. EcoCyc: fusing model organism databases with systems biology. Nucleic acids research. 2013;41(Database issue):D605-612.

87. Anjem A, Varghese S, Imlay JA. Manganese import is a key element of the OxyR response to hydrogen peroxide in Escherichia coli. Mol Microbiol. 2009;72(4):844–858.

88. Park SJ, Gunsalus RP. Oxygen, iron, carbon, and superoxide control of the fumarase fumA and fumC genes of Escherichia coli: role of the arcA, fnr, and soxR gene products. J Bacteriol. 1995;177(21):6255–6262.

89. Varghese S, Tang Y, Imlay JA. Contrasting sensitivities of Escherichia coli aconitases A and B to oxidation and iron depletion. J Bacteriol. 2003;185(1):221–230.

90. Eser M, Masip L, Kadokura H, Georgiou G, Beckwith J. Disulfide bond formation by exported glutaredoxin indicates glutathione’s presence in the E. coli periplasm. Proc Natl Acad Sci U S A. 2009; 106(5): 1572–1577.

91. Suzuki H, Koyanagi T, Izuka S, Onishi A, Kumagai H. The yliA, -B, -C, and -D genes of Escherichia coli K-12 encode a novel glutathione importer with an ATP-binding cassette. J Bacteriol. 2005;187(17):5861–5867.

92. Vuilleumier S. Bacterial glutathione S-transferases: what are they good for? J Bacteriol. 1997; 179(5): 1431–1441.

93. Summers AO, Wireman J, Vimy MJ, Lorscheider FL, Marshall B, Levy SB, Bennett S, Billard L. Mercury released from dental “silver” fillings provokes an increase in mercury-and antibiotic-resistant bacteria in oral and intestinal floras of primates. Antimicrob Agents Chemother. 1993;37(4):825–834.

94. US - Food & Drug Administration (FDA): Mercury Levels in Commercial Fish and Shellfish (1990–2012) http://www.fda.gov/Food/FoodborneIllnessContaminants/Metals/ucm115644.htm

95. McElwee MK, Freedman JH. Comparative toxicology of mercurials in Caenorhabditis elegans. Environ Toxicol Chem. 2011;30(9):2135–2141.

96. Zheng M, Wang X, Templeton LJ, Smulski DR, LaRossa RA, Storz G. DNA microarray-mediated transcriptional profiling of the Escherichia coli response to hydrogen peroxide. J Bacteriol. 2001;183(15):4562–4570.

97. Mehta HH, Liu Y, Zhang MQ, Spiro S. Genome-wide analysis of the response to nitric oxide in uropathogenic Escherichia coli CFT073. Microb Genom. 2015;1(4):e000031.

98. Rau MH, Calero P, Lennen RM, Long KS, Nielsen AT. Genome-wide Escherichia coli stress response and improved tolerance towards industrially relevant chemicals. Microb Cell Fact. 2016;15(1):176.

99. Yung PY, Grasso LL, Mohidin AF, Acerbi E, Hinks J, Seviour T, Marsili E, Lauro FM. Global transcriptomic responses of Escherichia coli K-12 to volatile organic compounds. Sci Rep. 2016;6:19899.

100. Wong GT, Bonocora RP, Schep AN, Beeler SM, Lee Fong AJ, Shull LM, Batachari LE, Dillon M, Evans C, Becker CJ et al. Genome-Wide Transcriptional Response to Varying RpoS Levels in Escherichia coli K-12. J Bacteriol. 2017;199(7).

101. Ung CY, Lam SH, Hlaing MM, Winata CL, Korzh S, Mathavan S, Gong Z. Mercury-induced hepatotoxicity in zebrafish: in vivo mechanistic insights from transcriptome analysis, phenotype anchoring and targeted gene expression validation. BMC Genomics. 2010;11:212.

102. Lash LH, Zalups RK. Alterations in renal cellular glutathione metabolism after in vivo administration of a subtoxic dose of mercuric chloride. J Biochem Toxicol. 1996; 11(1):1–9.

103. Lu X, Xiang Y, Yang G, Zhang L, Wang H, Zhong S. Transcriptomic characterization of zebrafish larvae in response to mercury exposure. Comp Biochem Physiol C Toxicol Pharmacol. 2017;192:40–49.

104. Kanai T, Takahashi K, Inoue H. Three distinct-type glutathione S-transferases from Escherichia coli important for defense against oxidative stress. J Biochem. 2006;140(5):703–711.

105. Xu FF, Imlay JA. Silver(I), mercury(II), cadmium(II), and zinc(II) target exposed enzymic iron-sulfur clusters when they toxify Escherichia coli. Appl Environ Microbiol. 2012;78(10):3614–3621.

106. Varghese S, Wu A, Park S, Imlay KR, Imlay JA. Submicromolar hydrogen peroxide disrupts the ability of Fur protein to control free-iron levels in Escherichia coli. Mol Microbiol. 2007;64(3):822–830.

107. Bagg A, Neilands JB. Ferric uptake regulation protein acts as a repressor, employing iron (II) as a cofactor to bind the operator of an iron transport operon in Escherichia coli. Biochemistry. 1987;26(17):5471–5477.

108. de Lorenzo V, Wee S, Herrero M, Neilands JB. Operator sequences of the aerobactin operon of plasmid ColV-K30 binding the ferric uptake regulation (fur) repressor. J Bacteriol. 1987;169(6):2624–2630.

109. Fontecave M, Choudens SO, Py B, Barras F. Mechanisms of iron-sulfur cluster assembly: the SUF machinery. J Biol Inorg Chem. 2005;10(7):713–721.

110. Zhao G, Ceci P, Ilari A, Giangiacomo L, Laue TM, Chiancone E, Chasteen ND. Iron and hydrogen peroxide detoxification properties of DNA-binding protein from starved cells. A ferritin-like DNA-binding protein of Escherichia coli. The Journal of biological chemistry. 2002;277(31):27689–27696.

111. Lund BO, Miller DM, Woods JS. Mercury-induced H2O2 production and lipid peroxidation in vitro in rat kidney mitochondria. Biochem Pharmacol. 1991;42 Suppl:S181–187.

112. Liochev SI, Fridovich I. Modulation of the fumarases of Escherichia coli in response to oxidative stress. Arch Biochem Biophys. 1993;301(2):379–384.

113. Martin JE, Imlay JA. The alternative aerobic ribonucleotide reductase of Escherichia coli, NrdEF, is a manganese-dependent enzyme that enables cell replication during periods of iron starvation. Mol Microbiol. 2011;80(2):319–334.

114. Jordan A, Aslund F, Pontis E, Reichard P, Holmgren A. Characterization of Escherichia coli NrdH. A glutaredoxin-like protein with a thioredoxin-like activity profile. The Journal of biological chemistry. 1997;272(29):18044–18050.

115. Sharma SK, Goloubinoff P, Christen P. Heavy metal ions are potent inhibitors of protein folding. Biochem Biophys Res Commun. 2008;372(2):341–345.

116. Arsene F, Tomoyasu T, Bukau B. The heat shock response of Escherichia coli. Int J Food Microbiol. 2000;55(1-3):3–9.

117. Bissonnette SA, Rivera-Rivera I, Sauer RT, Baker TA. The IbpA and IbpB small heat-shock proteins are substrates of the AAA+ Lon protease. Mol Microbiol. 2010;75(6): 1539–1549.

118. Gaubig LC, Waldminghaus T, Narberhaus F. Multiple layers of control govern expression of the Escherichia coli ibpAB heat-shock operon. Microbiology. 2011; 157(Pt 1):66–76.

119. Matuszewska E, Kwiatkowska J, Kuczynska-Wisnik D, Laskowska E. Escherichia coli heat-shock proteins IbpA/B are involved in resistance to oxidative stress induced by copper. Microbiology. 2008;154(Pt 6):1739–1747.

120. Wang A, Crowley DE. Global gene expression responses to cadmium toxicity in Escherichia coli. J Bacteriol. 2005;187(9):3259–3266.

121. Salmon K, Hung SP, Mekjian K, Baldi P, Hatfield GW, Gunsalus RP. Global gene expression profiling in Escherichia coli K12. The effects of oxygen availability and FNR. The Journal of biological chemistry. 2003;278(32):29837–29855.

122. Bibikov SI, Biran R, Rudd KE, Parkinson JS. A signal transducer for aerotaxis in Escherichia coli. J Bacteriol. 1997;179(12):4075–4079.

123. Pruss BM, Campbell JW, Van Dyk TK, Zhu C, Kogan Y, Matsumura P. FlhD/FlhC is a regulator of anaerobic respiration and the Entner-Doudoroff pathway through induction of the methyl-accepting chemotaxis protein Aer. J Bacteriol. 2003; 185(2):534–543.

124. Funk AE, Day FA, Brady FO. Displacement of zinc and copper from copper-induced metallothionein by cadmium and by mercury: in vivo and ex vivo studies. Comp Biochem Physiol C. 1987;86(1): 1–6.

125. O’Connor TR, Graves RJ, de Murcia G, Castaing B, Laval J. Fpg protein of Escherichia coli is a zinc finger protein whose cysteine residues have a structural and/or functional role. The Journal of biological chemistry. 1993;268(12):9063–9070.

126. Binet MR, Poole RK. Cd(II), Pb(II) and Zn(II) ions regulate expression of the metal-transporting P-type ATPase ZntA in Escherichia coli. FEBS Lett. 2000;473(1):67–70.

127. Petit-Hartlein I, Rome K, de Rosny E, Molton F, Duboc C, Gueguen E, Rodrigue A, Coves J. Biophysical and physiological characterization of ZraP from Escherichia coli, the periplasmic accessory protein of the atypical ZraSR two-component system. Biochem J. 2015;472(2):205–216.

128. Colaco HG, Santo PE, Matias PM, Bandeiras TM, Vicente JB. Roles of Escherichia coli ZinT in cobalt, mercury and cadmium resistance and structural insights into the metal binding mechanism. Metallomics. 2016;8(3):327–336.

129. Owens RA, Hartman PE. Export of glutathione by some widely used Salmonella typhimurium and Escherichia coli strains. J Bacteriol. 1986; 168(1): 109–114.

130. Smirnova GV, Muzyka NG, Oktyabrsky ON. Effects of cystine and hydrogen peroxide on glutathione status and expression of antioxidant genes in Escherichia coli. Biochemistry (Mosc). 2005;70(8):926–934.

131. Fitzgerald DM, Bonocora RP, Wade JT. Comprehensive mapping of the Escherichia coli flagellar regulatory network. PLoS Genet. 2014;10(10):e1004649.

132. Dong T, Yu R, Schellhorn H. Antagonistic regulation of motility and transcriptome expression by RpoN and RpoS in Escherichia coli. Mol Microbiol. 2011; 79(2):375–386.

133. De Lay N, Gottesman S. A complex network of small non-coding RNAs regulate motility in Escherichia coli. Mol Microbiol. 2012;86(3):524–538.

134. Mermod M, Magnani D, Solioz M, Stoyanov JV. The copper-inducible ComR (YcfQ) repressor regulates expression of ComC (YcfR), which affects copper permeability of the outer membrane of Escherichia coli. Biometals. 2012;25(1):33–43.

135. Schwechheimer C, Kuehn MJ. Outer-membrane vesicles from Gram-negative bacteria: biogenesis and functions. Nat Rev Microbiol. 2015;13(10):605–619.

136. Crepin S, Chekabab SM, Le Bihan G, Bertrand N, Dozois CM, Harel J. The Pho regulon and the pathogenesis of Escherichia coli. Vet Microbiol. 2011; 153(1-2):82–88.

137. Hsieh YJ, Wanner BL. Global regulation by the seven-component Pi signaling system. Curr Opin Microbiol. 2010;13(2):198–203.

138. Hondorp ER, Matthews RG. Oxidative stress inactivates cobalamin-independent methionine synthase (MetE) in Escherichia coli. PLoS Biol. 2004;2(11):e336.

139. Brooks P, Davidson N. Mercury(II) Complexes of Imidazole and Histidine. Journal of the American Chemical Society. 1960;82(9):2118–2123.

140. Pal C, Bengtsson-Palme J, Kristiansson E, Larsson DG. Co-occurrence of resistance genes to antibiotics, biocides and metals reveals novel insights into their co-selection potential. BMC Genomics. 2015;16(1):964.

141. Wireman J, Liebert CA, Smith T, Summers AO. Association of mercury resistance with antibiotic resistance in the gram-negative fecal bacteria of primates. Appl Environ Microbiol. 1997;63(11):4494–4503.

142. Mehta P, Casjens S, Krishnaswamy S. Analysis of the lambdoid prophage element e14 in the E. coli K-12 genome. BMC Microbiol. 2004;4:4.

143. Wang X, Kim Y, Ma Q, Hong SH, Pokusaeva K, Sturino JM, Wood TK. Cryptic prophages help bacteria cope with adverse environments. Nat Commun. 2010;1:147.

